# Dissecting the coordinated progression of cell states in spatial transcriptomics with CoPro

**DOI:** 10.64898/2026.04.17.719309

**Authors:** Zhen Miao, Yilong Qu, Sijia Huang, Linshan Laux, Samuel Peters, Alisha Aristel, Zhaojun Zhang, Laura Niedernhofer, Andrew McMahon, Junhyong Kim, Nancy R. Zhang

## Abstract

Spatial transcriptomics enables the study of how cells coordinate their molecular states within tissue, providing insight into both normal function and disease processes. A key challenge is to identify gene expression programs that vary continuously across space and are coordinated between cell types. We present **CoPro**, a computational framework for detecting the spatially coordinated progression of cellular states. CoPro can operate in both supervised and unsupervised modes to identify gene programs that co-vary within or between cell types, and to disentangle multiple overlapping spatial patterns. CoPro can be applied to single-cell-level spatial transcriptomics datasets, including MERFISH, SeqFISH+, Xenium, and histology-imputed transcriptomic data. We demonstrate the utility of CoPro with data collected from colon, brain, liver, and kidney tissues. In the colon, CoPro separates epithelial differentiation along the crypt axis from spatially localized inflammatory signals. In the aging liver, it identifies multiple aging-associated cellular programs superimposed on anatomical zonation. In the brain, the flexible kernel design enables the decoupling of the gene expression gradient along the dorsal-ventral and medial-lateral axes. In the kidney, CoPro identifies tubule-vasculature coordination that is essential in nephron function. These results demonstrate CoPro’s utility for analyzing spatial coordination of gene expression in complex tissues and disentangling overlapping biological processes, such as anatomical organization and disease-associated variation.

## Introduction

Tissue function emerges from spatially coordinated interactions among neighboring cells, which shape one another’s molecular state through direct contact, secreted signals, and shared microenvironments. These interactions are not only fundamental to maintaining homeostasis, they also drive pathological processes. Spatial transcriptomics^1^ offers an opportunity to study these interactions in situ, capturing the organization of cells into domains and microstructures that collectively determine tissue function and dysfunction.

Most computational methods for analyzing spatial transcriptomics data focus on segmenting tissues into discrete spatial domains, often referred to as spatial niches^2^. These domains are typically defined based on gene expression patterns^3^, cell type composition^4–6^, cell-cell proximity^7^, or multimodal features^8^. While this approach has been useful for initial data visualization and exploration, it may impose artificial categorization on what might be continuous spatial transitions. In this work, we show that, in many cases, discrete domains can be better described by the superposition of continuous axes of coordinated variation across cell types, such as differentiation gradients, metabolic zonation, and inflammatory responses, that can coexist within the same spatial neighborhood. We develop a framework to dissect these overlapping axes in a cell type-specific manner, resolving how distinct gene programs in different cell types co-progress along shared spatial gradients, with or without prior biological guidance.

Many recent methods have begun to address continuous spatial variation. These include MERINGUE^9^, DIALOGUE^10^, SpaceFlow^11^, ONTraC^12^, and GASTON^13^. However, these methods do not address the problem of identifying continuous, multi-axis spatial coordination of gene programs across multiple cell types at single-cell resolution. There remains a need for a computational framework that simultaneously preserves continuous single-cell spatial resolution, explicitly maps the coupled molecular progression of diverse cell populations, and disentangles multiple co-existing axes of spatial coordination.^10–13^

Another key analytical goal of spatial transcriptomics is the integrative analysis of multiple samples with spatial information. Comparing patterns across samples, whether it is for the validation of findings or the elucidation of cross-sample heterogeneity, requires a principled approach to multiple-sample integration. However, most existing methods are designed for single-sample analysis and do not explicitly support multi-sample settings^14,15^. Among methods that do accommodate multiple samples, many rely on spatial registration, assuming that tissue regions with similar histological appearance correspond to biologically comparable states^16–18^. CoPro relaxed this assumption and adopted an alternative approach by projecting all cells from different samples onto a biologically defined axis derived from shared gene expression programs. This approach enables interpretable cross-sample integration and reproducibility assessment in a manner that is robust to tissue morphology variation. We demonstrate the application with two examples, the first in a study of colon injury and the second in a study of liver aging.

CoPro explicitly models and detects continuous coordinated progression between spatially proximal cells. CoPro can operate in both supervised and unsupervised modes. In its supervised form, the user specifies a gene program of interest in one cell type, and CoPro identifies gene programs in other cell types that spatially co-vary with it. In unsupervised mode, CoPro discovers axes of coordinated variation de novo across cell types. We demonstrate supervised CoPro in the kidney tissue, whereas unsupervised CoPro is used for all other tissues.

## Results

### Overview of CoPro framework

We assume as input a spatial transcriptomics dataset in which each cell or pixel has been assigned to a broad cell type, based on transcriptomic clustering, morphological features, or external annotations (Fig. 1a). CoPro then models the coordinated variation of molecular programs between these cell types across space, with the goal of identifying cell type-specific gene expression patterns that progress jointly between spatially proximal cells.

**Figure 1.**
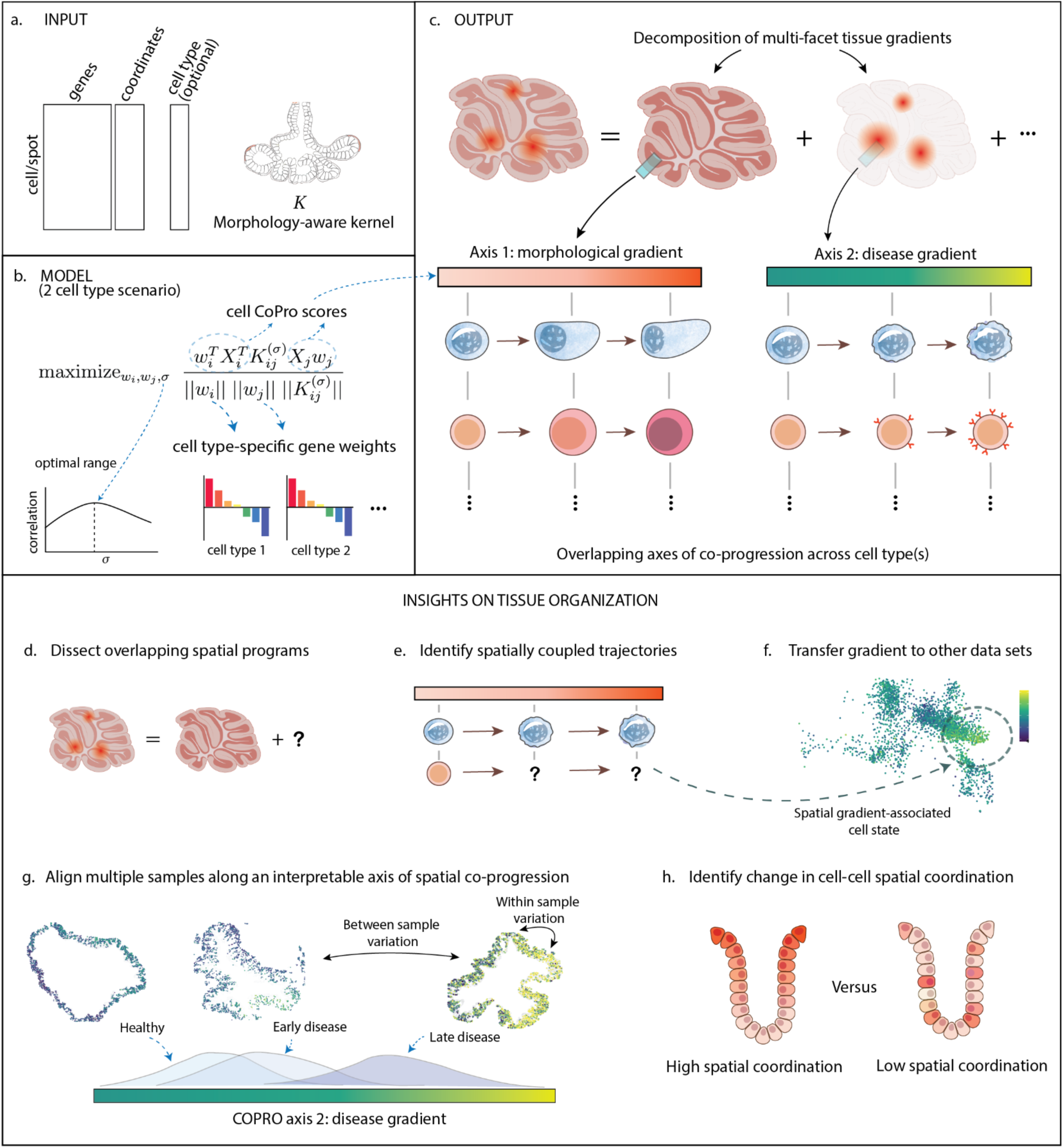
Overview of the CoPro computational framework. (**a**) Inputs to CoPro include cell-by-gene expression matrices, spatial coordinates, and optional cell type annotations, which are integrated through a spatial kernel. (**b**) CoPro identifies cell-type-specific gene weights by maximizing spatially weighted correlations between cell progression scores across cell types, with kernel bandwidth selected to capture the relevant spatial scale. (**c**) Decomposition of complex tissue organization into multiple orthogonal axes of spatial co-progression, each corresponding to a distinct biological process (for example, morphological or disease-associated gradients). (**d**) CoPro disentangles overlapping spatial programs within the same tissue into interpretable, non-redundant components. (**e**) In supervised mode, a known or inferred trajectory in one cell type can be used to identify spatially coupled trajectories in other cell types. (**f**) Learned gene weights can be transferred across datasets to map shared spatial gradients and associated cell states. (**g**) CoPro enables alignment of samples along a common axis of spatial co-progression, facilitating comparison across conditions such as disease progression. (**h**) Changes in the strength of spatial coordination between cell types can be quantified along each axis.

Even though CoPro can be applied to pixel-level data, as we demonstrate later, to describe the model, we will refer to each unit of observation as a “cell”. Suppose that we have chosen *T* cell types for co-progression analysis. For each cell type *t* ∈ {1,2, …, *T*}, let *n*_*t*_ be the number of cells assigned to the given type. We recommend first performing dimension reduction through PCA whitening to obtain a (scaled) *n*_*t*_ × *d*_*t*_ principal component matrix *X*_*t*_, where *d*_*t*_is the number of PCA components retained, capturing the expression variation within that type. Between every two cell types *s*, *t* we define a spatial kernel 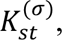 of dimensions *n*_*s*_ × *n*_*t*_, where the weight 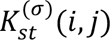 between every pair of cells *i*, *j*, with *i* of type *s* and *j* of type *t*, encodes their spatial proximity (**Methods**). *σ* denotes the bandwidth of cell-cell interaction.

Starting with the simplest case of two cell types and 1 sample (**Fig. 1b**), CoPro estimates canonical directions *w*_1_ ∈ ℜ^*d*1^ and *w*_2_ ∈ ℜ^*d*2^ to maximize 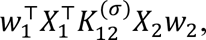 subject to the constraint that ‖*w*_*s*_‖ = 1 for *s* = 1,2. Thus, CoPro identifies combinations in the reduced feature space, specific to cell types 1 and 2, that maximize the kernelized correlation between them. Note that we could translate the weights in the reduced feature space to weights in gene space, via decoding the PCA embedding. In the general case of *T* ≥ 2 cell types, we would estimate *T* weight vectors *w*_1_, …, *w*_*T*_ by

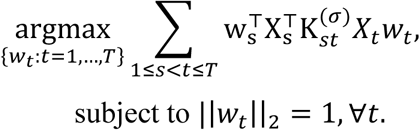

Finally, in the case of *q* ∈ {1,2, …, *Q*} samples, CoPro then solves the following optimization problem:

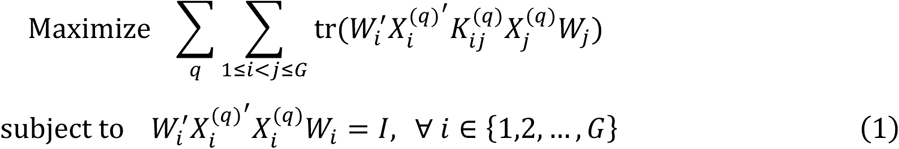

where *W*_*i*_ = [*w*_*i*,1_, *w*_*i*,2_, …, *w*_*i*,*K*_] is the canonical weight matrix for cell type *i*, with columns *w*_*i*,*k*_ representing a canonical weight vector to be estimated from data. The product 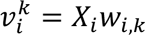 is termed the “cell progression score” for the *k*^th^ canonical component, representing an inferred spatial trend along the given spatial co-progression axis (**Fig. 1c**). This formulation identifies linear combinations of gene expression that are maximally spatially correlated across neighboring cells as defined by the kernel. The estimated weights *W*_*i*_ for each cell type reveal interpretable gene programs associated with the coordinated progression. The formulation is a generalization of the classical Canonical Correlation Analysis (CCA)^19^ by incorporating a spatial kernel and allowing for multiple (>2) interacting entities.

This formulation is distinct from Kernelized CCA (KCCA)^20–22^, which applies kernel functions to the gene expression space itself— implicitly mapping the data to a high-dimensional feature space in order to capture non-linear correlations between two omics modalities. In CoPro, the kernel operates on the spatial coordinate space: it encodes the proximity structure of the tissue microenvironment and re-weights the cross-cell-type covariance by the spatial neighborhood, while gene expression remains in the original (reduced-dimensional) embedding. In other words, CoPro introduces spatial context into a classical linear CCA objective.

To quantify the spatial co-progression across or within cell types, we define the Bidirectional Spatial Correlation (BSC) to capture the bidirectional spatial associations between cell progression scores. For two cell types with cell progression scores 𝑣_*i*_ and 𝑣_j_, the BSC is computed as:

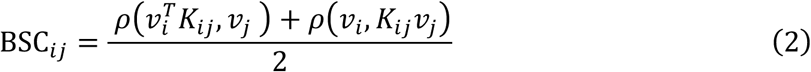

where *ρ* denotes Pearson correlation. The first term measures how the spatially-contextualized cell score of type *i* correlates with the cell score of type *j*, while the second term captures the reverse relationship. This bidirectional formulation ensures symmetry and provides an interpretable metric bounded in [-1, 1], with a similar interpretation as conventional correlation. Unlike traditional correlation measures that ignore spatial context and require matched number and index-level pairing of cells of each type, the BSC explicitly incorporates spatial proximity as represented by the kernel matrix, enabling detection of spatially coordinated biological programs between cell types. This metric thus provides a quantification of how cellular states progress in spatial coordination, revealing potential cell-cell communication and shared microenvironmental influences that drive tissue organization and function. We also define a kernel-normalized correlation for bandwidth (*σ*) selection, which has favorable statistical properties but is less directly interpretable (**Methods**).

CoPro dissects spatial transcriptomic data into overlapping spatial gradients, enabling the separation of distinct but coexisting spatial programs within the same tissue (**Fig. 1d**). It supports both unsupervised and supervised modes. In the unsupervised setting, it identifies axes of spatial coordination de novo, using only the expression matrix and the spatial position of cells. In the supervised setting, the user can provide a known or inferred spatial trend for one cell type, and CoPro will identify gene programs in other cell types that are spatially aligned with it, thereby uncovering spatially coupled cellular programs across types (**Fig. 1e**). These complementary modes can also be combined; for example, using unsupervised discovery to identify candidate axes, followed by supervised refinement or transfer to new datasets (**Fig. 1f**).

For cell types with well-defined differentiation gradients, CoPro’s inferred progression scores often recapitulate known expression-driven programs. However, its primary advantage lies in leveraging spatial context to uncover coordinated cellular transitions that would otherwise remain hidden. In particular, CoPro detects subtle or ambiguous transitions that lack sufficient signal for non-spatial approaches, and recovers spatial trends in rare or plastic cell types by borrowing information from their spatial coupling with more abundant populations. This cross-cell-type “spatial borrowing” is especially valuable in settings such as targeted gene panels, where gradient-informative genes may be sparse, and enables coherent spatial trends to be identified and compared across datasets.

By default, CoPro uses a Gaussian kernel with a bandwidth σ (the scale parameter controlling the effective range of spatial interactions) chosen to maximize the mean first-component kernel-normalized correlation across cell-type pairs (Methods). In tissues with multiple overlapping spatial trends, CoPro iteratively identifies orthogonal axes of co-progression of decreasing correlation strength. This allows spatial transcriptomic data to be decomposed into a set of interpretable, non-redundant gradients. Each axis is defined by cell-type-specific gene weights and corresponding spatial progression scores, which can be visualized to reveal coordinated variation across the tissue. These axes provide a common coordinate system that can be used to align samples along biologically meaningful gradients, facilitating comparisons across conditions or disease states (**Fig. 1g**). In addition to identifying axes, CoPro quantifies the strength of spatial coordination between cell types along each axis, enabling the detection of changes in cell–cell spatial coupling across conditions (**Fig. 1h**).

Because the optimization in (1) may capture spurious correlations that arise by chance, we implement a permutation-based procedure to assess the statistical significance of each identified axis. Specifically, the tissue is partitioned into non-overlapping spatial patches (default side length equal to twice the kernel bandwidth σ), and for each cell type being tested, spatial bins are randomly shuffled and cells at each bin’s locations are replaced by same-type cells sampled from the mapped target bin, while spatial coordinates are held fixed. This patch-based design preserves within-cell-type spatial autocorrelation while breaking cross-cell-type spatial coupling, thereby generating an appropriate null distribution (**Methods**).

### Benchmark analyses via simulations

To place CoPro in the context of existing approaches, we compared its conceptual scope and capabilities to representative methods for spatial transcriptomic analysis (**Supplementary Table 1**). Domain clustering approaches identify discrete spatial niches; trajectory-based methods such as ONTraC infer continuous spatial ordering based on cell type composition; SpaceFlow identifies transcriptional gradients at the neighborhood level; and DIALOGUE identifies multi-cell-type programs but does not explicitly model spatial organization at single-cell resolution. CoPro addresses a distinct analytical need: the identification of spatially coupled continuous programs of co-progression across cell types. It resolves overlapping gradients rather than partitioning tissue into discrete regions, and incorporates spatial proximity directly into the optimization at single-cell resolution. Because these methods address different analytical objectives, direct benchmarking is inherently limited: strong performance by one method on a task it was designed for does not imply superiority over methods designed for different goals. Accordingly, in the simulation studies below, we compare CoPro against methods with the most overlapping goals, specifically, the ability to estimate continuous spatial cell scores—while using **Supplementary Table 1** to provide a broader conceptual comparison. We emphasize that benchmark results should be interpreted in light of each method’s design intent.

Here, we designed a series of simulation studies based on realistic gene expression profiles from the Liver Cell Atlas^23^. These simulations assess CoPro’s ability to recover known spatial patterns in scenarios of increasing complexity: two cell types with a single gradient, three cell types with varying proportions, and finally multiple orthogonal axes of tissue organization.

Our simulation framework generates spatial transcriptomics data by assigning cells from real single-cell RNA-seq datasets to spatial locations under controlled patterns of spatial organization (**Fig. 2a**; **Methods**). Rather than simulating gene expression de novo, this framework preserves realistic gene expression levels and gene–gene correlations by using observed single-cell profiles, which are matched to spatial positions according to specified spatial gradients. This design allows explicit control over whether and how cell types exhibit coordinated progression across space. In particular, we can simulate null scenarios in which spatial autocorrelation is present within individual cell types but not shared across types (i.e. the absence of co-progression), as well as alternative scenarios in which two or more cell types are spatially coupled and co-progress together. By preserving the statistical structure of real transcriptomic data while systematically varying spatial coordination patterns, this framework enables controlled evaluation of methods across a wide range of settings.

**Figure 2.**
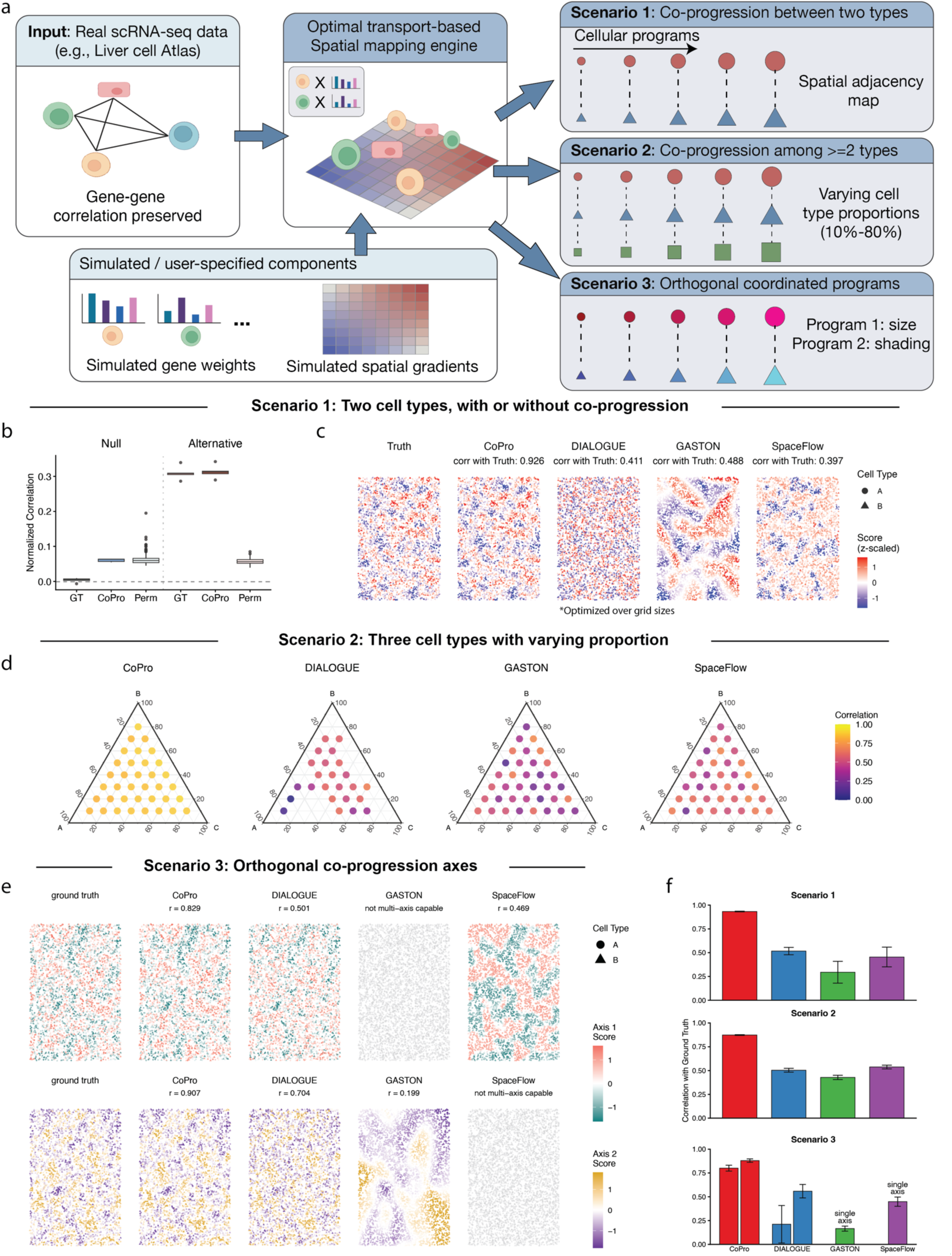
Systematic benchmarking of CoPro against state-of-the-art methods using realistic simulations. **(a)** Overview of the simulation framework. Real scRNA-seq data (e.g., Liver Cell Atlas) serves as input to preserve gene–gene correlations. An optimal transport-based engine maps cells to spatial locations based on user-specified components, generating three scenarios: (1) Co-progression between two cell types; (2) Co-progression among three cell types with varying proportions (10%–80%); and (3) Multiple orthogonal coordinated programs (e.g., size vs. shading). **(b)** Assessment of signal detection capability. Boxplots display the normalized correlation between cell types under Null (no co-progression) and Alternative (co-progression) conditions. CoPro estimates (middle) closely match the Ground Truth (GT, left) in both scenarios, distinguishing true signal from noise (Perm, right). **(c)** Qualitative comparison of spatial pattern recovery in Scenario 1 (two cell types). Spatial plots show ground truth scores versus scores estimated by CoPro, DIALOGUE, GASTON, and SpaceFlow. (**d**) Robustness to cell type composition in Scenario 2 (three cell types). Ternary plots visualize performance (color scale: correlation with ground truth) across all possible proportion combinations. CoPro maintains high accuracy regardless of cellular abundance, whereas DIALOGUE, GASTON, and SpaceFlow exhibit sensitivity to imbalances. (**e**) Recovery of orthogonal gradients in Scenario 3. Spatial plots show the first two axes of variation. CoPro accurately recovers distinct orthogonal programs as separate components. Other methods fail to separate the axes or capture only a single dimension. (**f**) Quantitative summary of performance. Bar plots show the mean Pearson correlation with ground truth across five independent runs for all scenarios. CoPro consistently outperforms DIALOGUE, GASTON, and SpaceFlow. Note that GASTON and SpaceFlow are not multi-axis capable in Scenario 3.

An important consideration for interpreting benchmark results is that each spatial simulation draws only a subset of cells from the full single-cell atlas, so the gene expression patterns in the spatial sample will inevitably deviate slightly from those in the original dataset. This sampling effect sets a natural ceiling on how well any method can recover the ground-truth gradient — even a perfect method operating on the sampled data cannot exceed the correlation between the gradient reconstructed from the sampled cells and the original gradient. We therefore benchmarked all methods against this theoretical performance ceiling rather than against the original gradient directly (**Methods**).

We first evaluated methods in the simplest scenario involving two cell types with a single coordinated spatial gradient. Using hepatocytes and endothelial cells, we simulated spatial patterns where both cell types exhibited correlated spatial organization reflecting a shared tissue gradient.

After applying Gaussian kernel smoothing to generate spatially coherent patterns, we observed clear spatial structure in the smoothed cell scores, with cells clustered by their position along the gradient axis (**Fig. 2a**). Of note, unlike many existing simulation frameworks that assume a strong global gradient (e.g. a linear, blocked, or other smooth change across the entire tissue), our simulation strategy produces more complex, spatially localized patterns that are harder to dissect.

In the two-cell-type simulations, we first evaluated CoPro’s performance with or without true coordinated progression between cells (**Fig. 2b**). Since CoPro always tries to find the optimal coordinated axis, its estimated correlation value is greater than zero even when there is no global co-progression. However, our patch-based permutation method yields the correct null distribution that successfully distinguishes real co-progression patterns from the null. When true coordinated progression exists, CoPro reliably recovers the underlying spatial gene expression gradient, even when there is no strong global gradient (**Fig. 2c**). In comparison, naïve permutation of spatial coordinates that do not account for within-cell-type autocorrelation fails to distinguish alternative from the null.

We then compared CoPro’s performance to DIALOGUE, SpaceFlow, ONTraC, and GASTON, which can each estimate a continuous spatial pattern (**Fig. 2c**). We note that DIALOGUE discretizes spatial coordinates into grids and identifies gene programs that are correlated across cell types within each grid cell. To ensure fair comparison, we optimized DIALOGUE’s grid size parameter across a range of values, selecting the configuration that maximized performance for each simulation. In SpaceFlow and GASTON, there is no global hyperparameter that governs the spatial bandwidth for the continuous cell score inference. CoPro substantially outperformed other methods across five independent simulation runs (mean Pearson r = 0.93, **Fig. 2c, Supplementary Table 2**). DIALOGUE, SpaceFlow, and GASTON recovered substantially less of the ground-truth gradient — DIALOGUE tending to resolve overly fine-grained patterns and SpaceFlow and GASTON tending toward coarser ones. These results highlight CoPro’s advantages in both accuracy and reliability for detecting spatially coordinated gene programs.

Real tissues contain multiple cell types in varying proportions, and the relative abundance of different cell types may affect method performance. To evaluate robustness to cell type composition, we extended our simulations to include three cell types and systematically varied their proportions using a ternary design (**Fig. 2d**). This design spans the full range of possible three-way proportions.

CoPro maintained high performance across all proportion configurations (mean Pearson r = 0.87) regardless of cell type composition (**Fig. 2d, Supplementary Table 3**). Performance was stable regardless of which cell type was most abundant, demonstrating that CoPro does not require balanced cell type compositions to detect coordinated spatial patterns. In contrast, existing methods showed lower and more variable performance across the same configurations, with correlations typically ranging from 0.4 to 0.6 and greater sensitivity to cell type proportions.

Complex tissues often exhibit multiple independent axes of organization. For example, the intestinal epithelium, as we will show later, has both a crypt-luminal axis reflecting cellular differentiation and inflammatory gradients overlayed onto this anatomical structure. To evaluate CoPro’s ability to dissect multiple overlapping spatial gradients, we simulated data with two orthogonal axes of coordinated gene expression (**Methods**). Notably, most existing methods are not designed to resolve multiple overlaid gradients: approaches such as GASTON and SpaceFlow typically recover a single dominant spatial manifold, while DIALOGUE can, in principle, capture multiple programs but relies on neighborhood binning and is therefore sensitive to changes in cell type composition.

CoPro successfully recovered both orthogonal axes across independent simulation runs (**Fig. 2f**). Importantly, CoPro correctly identified the two axes as separate canonical components rather than conflating them (mean Pearson r = 0.84 for each axis), demonstrating its ability to decompose complex tissue organization into distinct biological programs. Visual inspection of the spatial patterns confirmed that CoPro’s estimated cell scores recapitulated the spatial gradients present in the ground truth, with clear separation between the two axes (**Fig. 2e, Supplementary Table 4**).

These simulation studies establish CoPro as an accurate and robust method for detecting coordinated spatial gene expression patterns. The method’s stable performance across these conditions suggests it will be effective for analyzing real spatial transcriptomics datasets, where ground truth patterns are unknown and tissue composition varies across samples and conditions.

Even in settings dominated by a strong global spatial gradient, CoPro enables the dissection of multiple overlaid axes of organization. To illustrate and benchmark this capability, we applied CoPro to a real MERFISH spatial transcriptomics data studying the brain tissue organization (**Fig. 3a**). The original study reported that many cell types exhibit continuous expression dynamics along specific spatial axes, rather than forming distinct clusters. Here, we focus on the striatum to demonstrate CoPro’s distinctive capabilities. Within the striatum, both D1 and D2 medium spiny neurons display prominent gene expression gradients. The dominant spatial gradient is sufficiently strong that the first PC axis, computed without incorporating any spatial information, primarily recapitulates a dorsolateral-ventromedial pattern.

**Figure 3.**
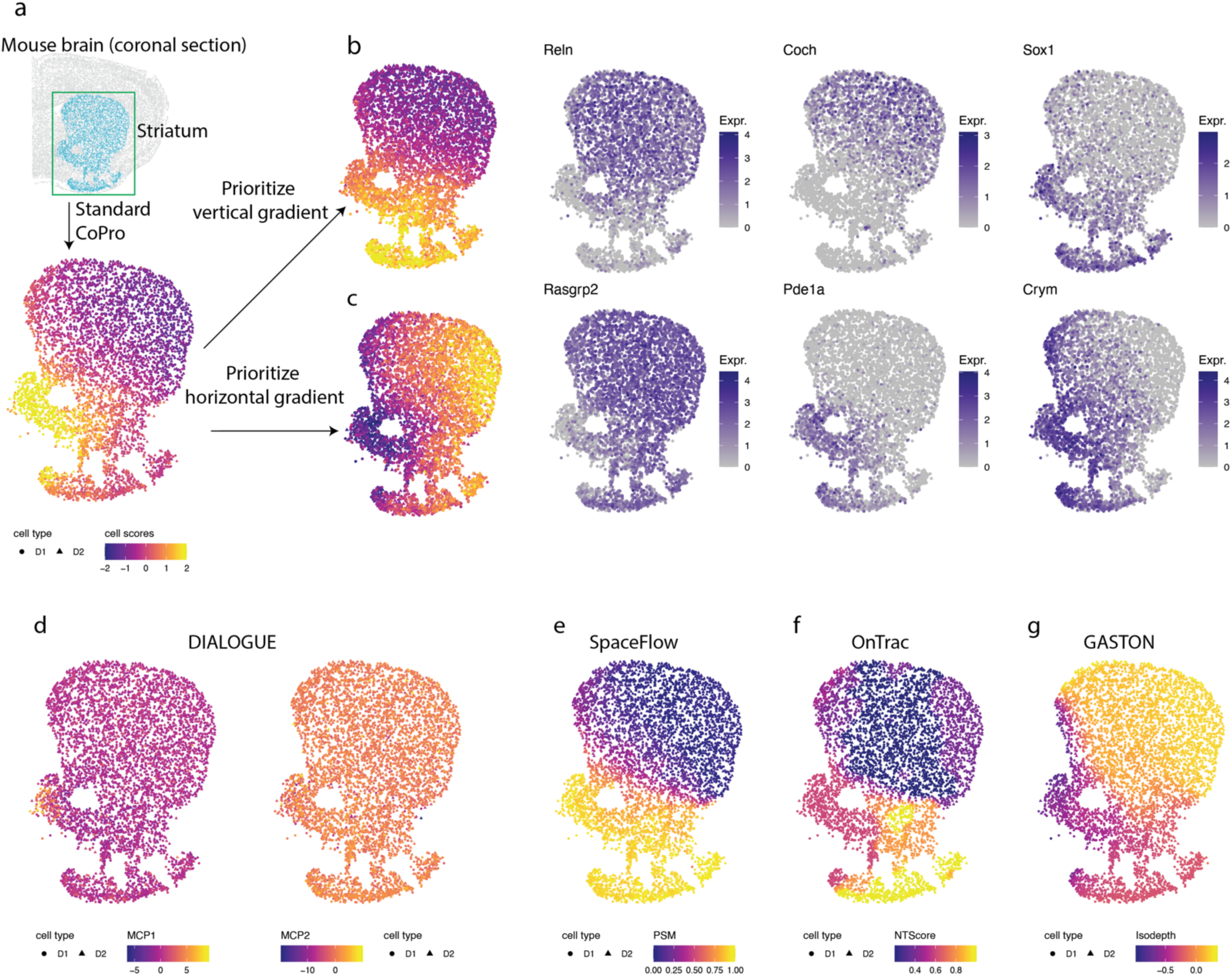
CoPro dissects orthogonal spatial axes in the mouse striatum. **(a)** CoPro analysis of D1 and D2 medium spiny neurons in the striatum region of a coronal brain section. **(b-c)** Left: By adjusting the spatial kernel to prioritize specific geometric axes, CoPro resolves two distinct, orthogonal gradients: a vertical (dorsal-ventral) axis (**b**) and a horizontal (lateral-medial) axis (**c**). Right: Example of top weighted gene associated with the vertical (top) and horizontal (bottom) axes. **(d-g)** Comparison with state-of-the-art methods: DIALOGUE (MCP1 and MCP2), SpaceFlow (PSM), OnTrac (NTScore), and GASTON (isodepth).

However, biological tissues often contain multiple, orthogonal gradients that are difficult to disentangle using standard dimensionality reduction. The flexibility of kernel selection in CoPro enabled us to overcome this by identifying spatial gradients along any user-specified axis. By applying different kernel configurations, we successfully decomposed the spatial variance to find distinct gradients along both the dorsal-ventral and lateral-medial axes (**Fig. 3b-c**, **Methods**). Along the dorsal-ventral axis, CoPro identified *Reln*, *Coch*, and *Sox1* as top-weighted genes, consistent with previous results from single cell studies^24^. For the lateral-medial axis, CoPro highlighted *Rasgrp2*, *Pde1a*, and *Crym* as top-weighted genes, revealing orthogonal, axis-specific molecular signatures that characterize brain tissue organization.

Finally, we benchmarked CoPro against other spatial domain and gradient identification methods (**Fig. 3d-g**). We found that existing methods struggled to decompose the smooth, multi-directional continuous dynamics present in the striatum. SpaceFlow, OnTrac, and GASTON all prioritized the dorsal-ventral axis, and the first axis in DIALOGUE highlighted the nucleus accumbens from other regions. Crucially, none of these methods possesses the flexibility to disentangle the superimposed spatial gradients (dorsal-lateral and ventral-medial), demonstrating CoPro’s superior utility in resolving complex, overlapping spatial patterns.

### Colon injury and repair: dissecting morphological and inflammatory gradients

To illustrate how CoPro resolves multiple, biologically distinct axes of spatial organization in a dynamic tissue context, we reanalyzed a MERFISH spatial transcriptomics dataset of the mouse colon spanning the course of acute injury to peak inflammation and recovery (Day 0, 3, 9, and 21). The dataset includes a panel of 940 genes. The original study identified ten mucosal “cellular neighborhoods” representing distinct niches, ranging from healthy (MU1-3), acute injury (MU4), inflammatory (MU6-8), and repairing (MU9-11) states^6^. We used CoPro to interrogate how spatially coordinated gene programs shift over time, focusing first on early-stage injury (Days 0 and 3).

In the Day 0 samples, the colon epithelium displays a well-defined crypt morphology, with Lgr5+ stem cells located at the base^25^ and differentiated cell types positioned toward the lumen. This spatially organized differentiation process is tightly coupled with stromal and immune components, providing an ideal setting for evaluating CoPro’s ability to detect structured co-progression across cell types.

To start, we applied CoPro to epithelial cells, fibroblasts, and macrophages in Day 0 samples. By projecting the co-progression scores in situ (**Fig. 4a, Supplementary Fig. 1a**), we were able to recover the crypt-luminal morphological axis characterized by differentiation of epithelial cells from the crypt base (high scores) to the tip (low scores). This spatial variation is concordant with the low-dimensional embedding of the epithelia differentiation trajectory (**Fig. 4a**, middle panel). Comparison with the subtype annotations from the original publication confirmed that CoPro correctly ordered cells by differentiation stage, with stem cells at the base, colonocytes at the tip, and transit amplifying (TA) cells in between.

**Figure 4.**
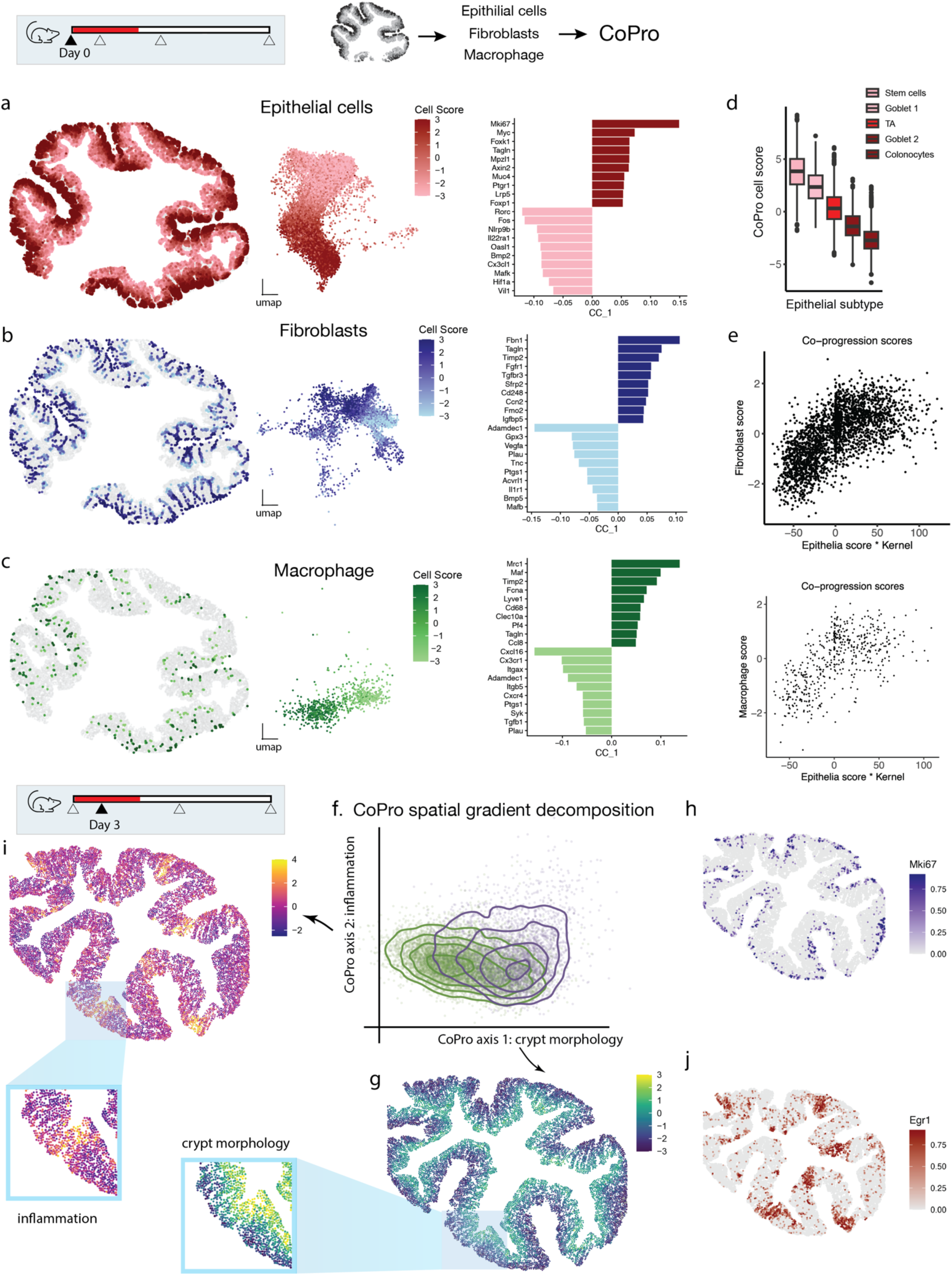
CoPro dissects morphological and inflammatory gradients in healthy and injured colon. (**a–c**) Application of CoPro to Day 0 mouse colon spatial transcriptomics for (**a**) Epithelial cells, (**b**) Fibroblasts, and (**c**) Macrophages. For each cell type, panels display: (Left) Spatial projection of CoPro scores recovering the crypt base-to-tip gradient; (Middle) UMAP embeddings colored by cell score; (Right) Bar plots of the top positive (crypt base/niche-support) and negative (crypt tip/differentiation) gene weights driving the co-progression. (**d**) Box plots of CoPro cell scores across epithelial subtypes defined in the original study. The scores recover the differentiation trajectory from Lgr5+ stem cells (crypt base) to differentiated colonocytes (crypt tip). (**e**) Scatter plots showing the correlation of Fibroblast (top) and Macrophage (bottom) co-progression scores against the Epithelial score, indicating spatially coordinated cellular states. (**f**) CoPro spatial gradient decomposition of Day 3 (acute injury) samples, resolving two orthogonal axes: Axis 1 (crypt morphology) and Axis 2 (inflammation). (**g**) Spatial projection of CoPro Axis 1, representing the underlying crypt morphology. (**h**) Spatial expression of Mki67, a proliferation marker enriched in the crypt base, corresponding to the crypt morphology axis (Axis 1). (**i**) Spatial projection of the combined CoPro axes. Insets highlight the distinct spatial distributions: the preserved continuous crypt morphology gradient (bottom) and the patchy, injury-responsive inflammation zones (top). (**j**) Spatial expression of Egr1, a representative immediate early gene (IEG) enriched in the patchy inflammation axis (Axis 2).

CoPro also identified corresponding co-varying states of fibroblast and macrophage (**Fig. 4b-c**). Fibroblast populations are known for their pronounced transcriptional plasticity, reflected here by the complexity of their low-dimensional embedding, which lacks a clear trajectory (**Fig. 4b**). Nonetheless, anchored by the strong epithelial gradient, CoPro was able to uncover a coherent co- varying transcriptomic axis within the fibroblast populations. In the absence of a consensus naming convention, the original study labeled fibroblast subtypes numerically. Based on marker gene expression, fibroblast 6 was linked by the original study to the crypt base, and fibroblast 2 to the tip (**Supplementary Fig. 1b**). This pattern was consistent with the reported cell scores from CoPro. The localization of Macrophage subpopulations had not been extensively characterized, but CoPro revealed that *Itgax+* macrophages localize to the tip and *Mrc1+* macrophages localize near the base (**Supplementary Fig. 1c**). The co-progression scores between any cell type pair were strongly correlated, confirming that all three cell types share a common spatial axis of coordinated progression along the crypt axis (**Fig. 4e**, **Supplementary Fig. 1d**).

CoPro provided interpretable gene weights that accurately captured the crypt base-to-tip gradient across the three cell types (**Fig. 4a-c**, right panels, **Supplementary Table 5**). Positive weights (crypt base-enriched) highlighted proliferation markers (*Mki67*, *Cenpf*) in epithelial cells, niche-supporting signaling (*Wnt2b*, *Fgfr1*) in fibroblasts, and tissue-supportive M2-like markers (*Mrc1*, *Cd163*) in macrophages. Conversely, negative weights (crypt tip-enriched) captured differentiation markers (*Bmp2*), antimicrobial programs (*Bmp5*), and immune activation genes (*Itgax*) associated with luminal surveillance. These gene weights were consistent with previous single cell studies^26,27^.

We next applied CoPro to Day 3 post-DSS treatment samples. Using a niche-based permutation test, we retained two significant spatial co-progression axes (**Supplementary Fig. 1e**). The first spatial axis (Axis 1) recapitulated the base-to-tip anatomical gradient (**Fig. 4f-g**), marked again by Mki67 expression at the base, suggesting that crypt morphology remains largely intact at this stage and continues to dominate as the principal co-progression axis (**Fig. 4g**). The second axis (Axis 2) revealed a patchy distribution enriched at the crypt tip (Fig. 4i, inset), corresponding to acute injury responses. Gene weights for Axis 2 highlighted coordinated upregulation of immediate early genes (IEGs^28,29^), such as *Egr1*, *Fosb*, *Fos*, and *Nr4a1*, across all cell types (**Fig. 4j**, **Supplementary Table 6**). Consistently, the high-scoring regions are enriched in mucosal microenvironment 4 identified in the original study (**Supplementary Fig. 1f-g**). However, whereas the original study characterized early stress response via a discrete microenvironment clustering, CoPro revealed it as a continuous directed co-progression linking spatially adjacent epithelial, fibroblast, and macrophage cells.

This orthogonal decomposition establishes CoPro’s ability to resolve concurrent biological processes – intact crypt-luminal morphology and acute stress responses – among co-localized cell types.

### Colon injury and repair: A unified axis of disease progression across samples

By Day 9 post-injury, the colons are in the severe disease stage, where the crypt morphology is largely disrupted. The original study noted substantial variation between mice and even between tissue sections from the same mice, making it difficult to summarize disease progression using discrete clustering, as many clusters become mouse- or sample-specific^6^. We sought to resolve this issue by using CoPro to obtain a holistic view of inter-sample disease variation relative to within-sample cellular variation.

We first applied CoPro to the Day 9 and Day 21 samples and captured the major tissue gradients associated with inflammation and repair, respectively (**Supplementary Fig. 2a-d**). This is evident given the gene programs aligned with each axis (**Supplementary Fig. 2e-f, Supplementary Table 8**) and their overlap with the manually curated neighborhood signatures identified by the original study (**Fig. 5a**).

**Figure 5.**
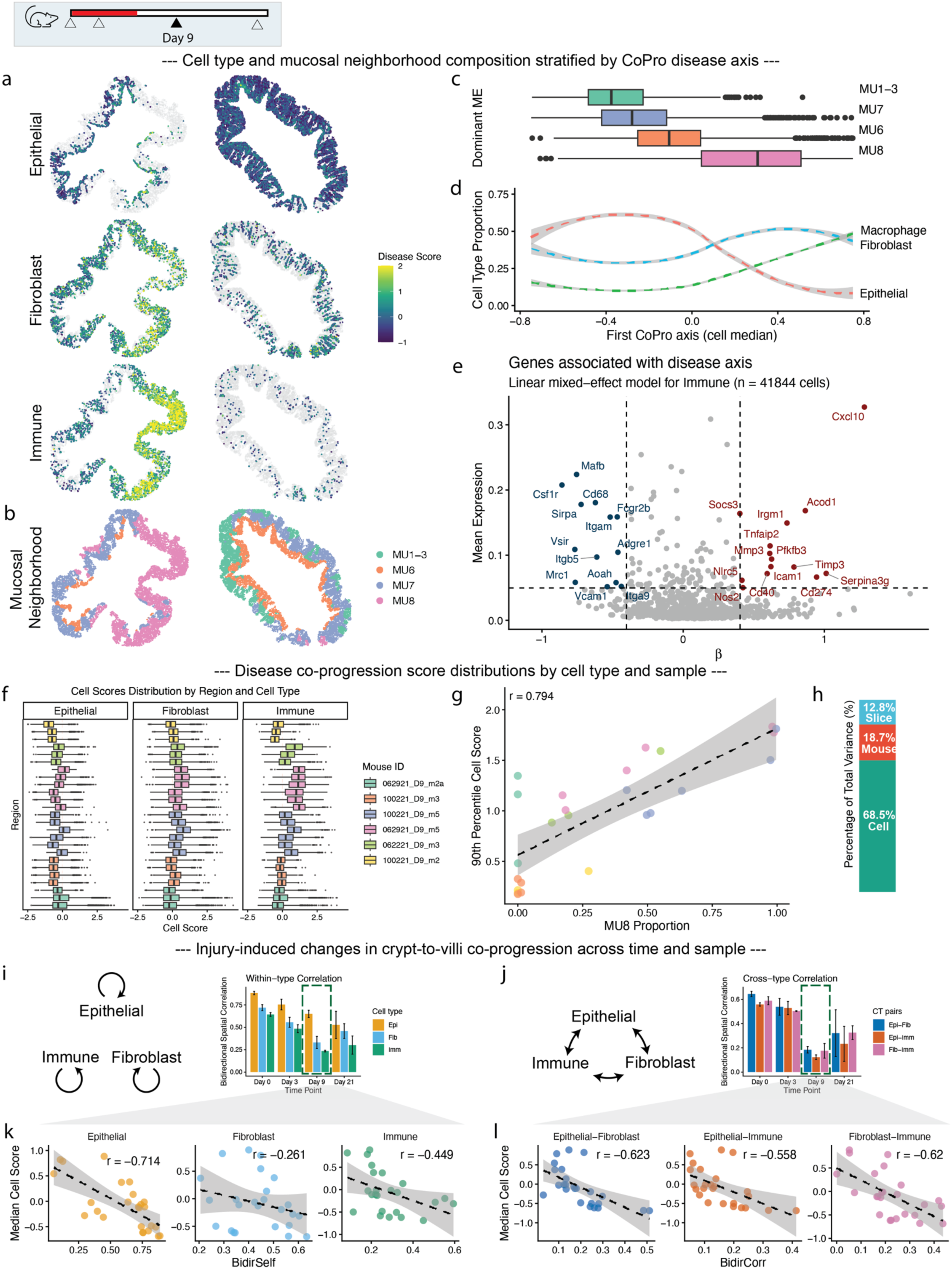
CoPro defines a unified axis of disease progression and quantifies the breakdown of tissue organization. (**a–e**) CoPro analysis of Day 9 colon samples identifies a unified disease axis. (**a–b**) Spatial projection of cell types (**a**) and Mucosal Neighborhoods (**b**) stratified by the CoPro disease axis. (**c**) Distribution of the first CoPro axis (cell median) across dominant microenvironments, showing the highest scores in the inflamed MU8 neighborhood. (**d**) Changes in cell type proportion along the first CoPro axis. (**e**) Genes associated with the disease axis in immune cells, identified using a linear mixed-effect model. Positive weights *β* indicate upregulation of inflammatory markers such as *Cxcl10*, *Socs3*, and *Mmp3* with increasing disease score. (**f–h**) Validation of the unified disease score across samples. (**f**) Cell score distributions stratified by region and cell type across different mouse replicates. (**g**) Scatter plot showing a strong correlation (0.774) between the 90th percentile CoPro cell score and the proportion of the MU8 neighborhood (inflamed niche) per sample. (**h**) Variance decomposition of disease scores, showing that cell-level variation (68.5%) dominates over mouse (18.7%) and slice (12.8%) effects. (**i–l**) Injury-induced changes in crypt-to-villi co-progression. (**i–j**) Bar plots of Bidirectional Spatial Correlation (BSC) computed for the homeostatic crypt-villus axis transferred to injured time points. Both within-type correlation (**i**) and cross-type correlation (**j**) drop sharply at Day 9. (**k–l**) Scatter plots showing the negative correlation between disease severity (Median Cell Score) and spatial coordination (BSC) for self-correlations (**k**) and cross-type pairs (**l**), indicating that disease progression disrupts both intra- and inter-cellular spatial organization.

To enable systematic cross-sample comparison, we first performed CoPro on a sample with mild disease to capture the full spectrum of healthy and diseased regions, designating the first CoPro axis as the disease progression axis. We then transferred the learned cell type-specific gene weights to all other samples, thereby computing a per-cell disease progression score. This approach enabled direct comparison of the distribution of disease progression scores across samples. The resulting scores successfully captured major tissue gradients associated with inflammation (**Fig. 5a**). The disease progression axis aligned strongly with the highly inflamed MU8 neighborhood (**Fig. 5b-c**) and revealed changes in cell type proportions associated with disease severity (**Fig. 5d**). Unlike discrete clustering, which can obscure transitions, CoPro scores mapped epithelial, fibroblast, and immune states onto a continuous manifold reflecting the degree of disease progression, linking spatially adjacent neighborhoods. We validated this unified axis by correlating the per-sample 90th percentile CoPro score with the abundance of the manually annotated MU8 neighborhood, yielding a strong correlation (R = 0.774) (**Fig. 5g**). This confirmed that CoPro captured disease severity in an unsupervised manner. Variance decomposition using a linear mixed-effects model revealed that cell-to-cell variation accounted for the largest share of variance (68.5%), indicating strong regional heterogeneity within tissues, while mouse-level effects (18.7%) exceeded section-level effects (12.8%) (**Fig. 5h**).

Using the unified disease axis, we identified cell-type-specific genes associated with disease severity, after adjusting for mouse- and section-level variation (**Fig. 5e**, **Supplementary Fig. 2g-h**, **Supplementary Table 7**). For example, in immune cells, higher disease scores were linked to the upregulation of inflammatory markers such as *Cxcl10*, *Socs3*, and *Mmp3,* recapitulating patterns observed in previous single cell studies^30,31^.

### Colon injury and repair: Characterizing the breakdown and recovery of crypt organization

Severe disease markedly disrupts the base-to-tip crypt axis that defines healthy colon organization. To quantify this disruption, we transferred the homeostatic crypt-luminal gene weights from Day 0 samples to Day 3, 9, and 21 samples. We then computed the Bidirectional Spatial Correlation (BSC), a metric quantifying the tightness of spatial coordination along the crypt–villus axis within and between cell types.

BSC values were high at Day 0 but diminished sharply at Day 9, reflecting the loss of structural organization (**Fig. 5i-j**). By Day 21, epithelial cells exhibited the greatest degree of structural restoration, whereas fibroblasts and immune cells remained largely disorganized (**Fig. 5i**). Notably, cross-type BSC values were consistently lower than within-type BSC (**Fig. 5j**), and both metrics showed a strong negative correlation with disease severity (**Fig. 5k, l**). These results indicate that disease progression involves not only the loss of epithelial differentiation but also the breakdown of spatial coordination between epithelial, stromal, and immune compartments. These observations suggest that full tissue recovery may involve re-establishing this intercellular alignment.

### Healthy liver: robust recovery of lobular zonation from histology-imputed spatial transcriptomics

The mammalian liver is organized into lobules, within which hepatocytes and non-parenchymal cells are arranged along a periportal–pericentral axis shaped by gradients of oxygen and nutrients. This organization underlies metabolic zonation and is maintained by coordinated signaling between hepatocytes and liver sinusoidal endothelial cells (LSECs)^32^. To further test and demonstrate CoPro, we asked whether it could recover this canonical spatial organization in a particularly challenging analytical setting: histology-imputed spatial transcriptomic data derived from H&E images integrated with the same-slide Visium assay, where expression is computationally inferred at 8-μm resolution by iSTAR^33^ (**Fig. 6a-b**). Here, individual pixels may capture partial or mixed cellular content, and technical artifacts from both histology and Visium-guided imputation are unavoidable. Following the terminology in iSTAR^33^, we call each 8-μm image patch a super-pixel.

**Figure 6.**
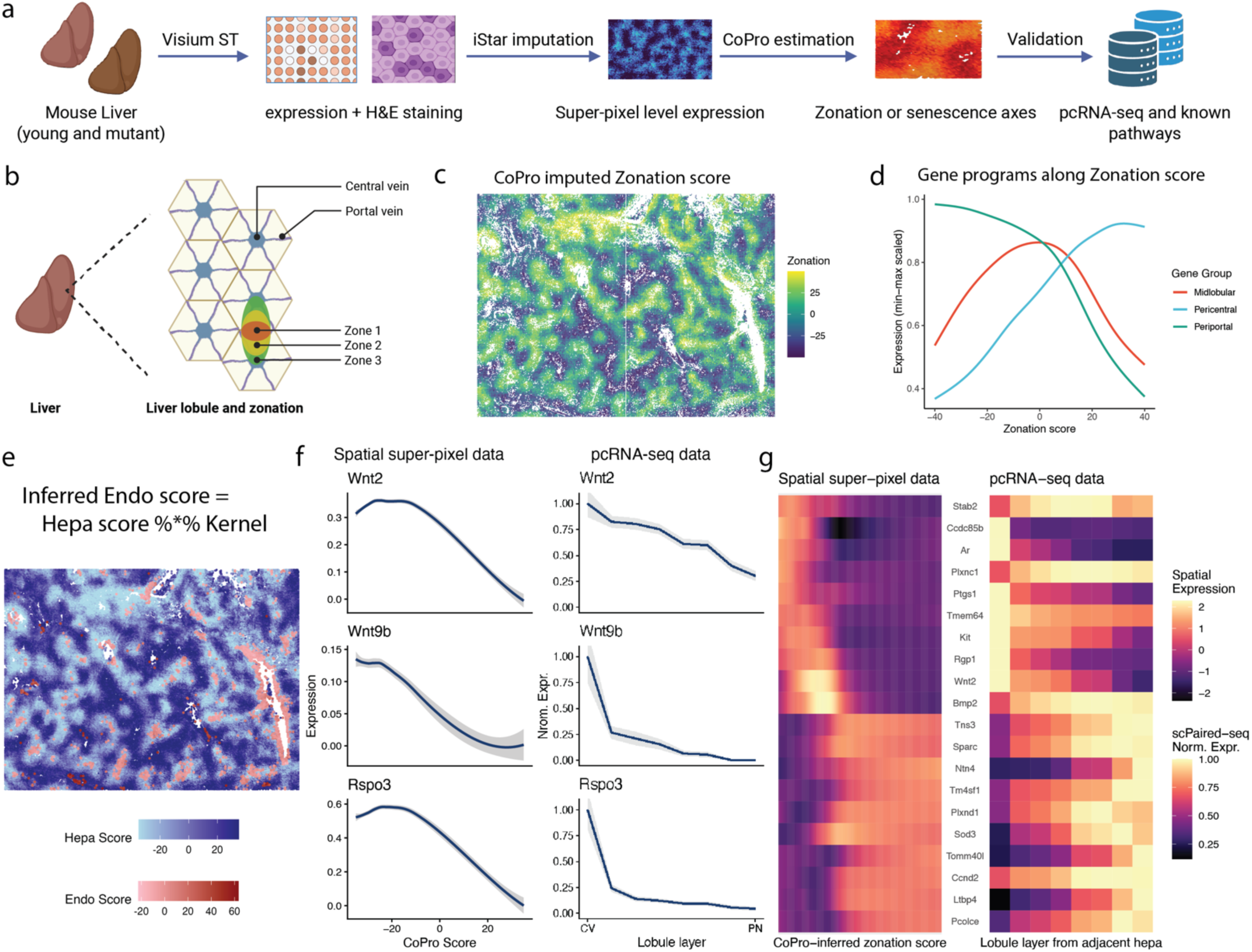
Dissection of liver lobule zones and hepatocyte-endothelial coordination using CoPro. (**a–b**) Overview of liver tissue organization and analysis workflow. (**a**) Visium spatial transcriptomics and H&E staining of wild-type (young) and *Ercc1*^-/Δ^ (mutant/aging) mouse livers. (**b**) Workflow: iStar super-resolution imputation is applied to H&E and expression data, followed by CoPro estimation to resolve zonation and senescence axes. (**c**) Spatial projection of CoPro-imputed zonation scores, recovering the hexagonal liver lobule structure with clear separation of Central Vein (Zone 3) and Portal Vein (Zone 1) regions. (**d**) Expression trends of gene groups along the inferred zonation score, delineating distinct Pericentral (red), Midlobular (green), and Periportal (blue) profiles. (**e–g**) Inference of endothelial cell zonation. (**e**) Projection of endothelial cell scores (Endo score) inferred from the hepatocyte-defined gradient (Hepa score) using the spatial kernel. (**f**) Heatmap comparing spatial super-pixel expression of top zonation genes with reference paired-cell RNA-seq (pcRNA-seq) data, showing strong concordance. (**g**) Spatial expression gradients of key endothelial morphogens (Wnt2, Wnt9b, Rspo3) plotted against CoPro score (left) and lobule layer (right). (**h**) Decomposition of the aging (*Ercc1*^-/Δ^) liver into four distinct transcriptional components. Left panels display spatial score projections for each component on tissue patches (C: Pericentral, M: Midlobular, P: Periportal).

We start with healthy wild-type mouse liver. Conventional clustering or trajectory-based approaches applied to either the original Visium spots or the imputed super-pixels resulted in a mixed cell cloud where the strong lobular zonation axis was obscured by technical noise and inherent mixed-cell data (**Supplementary Fig. 3a-i**). We therefore first applied CoPro in a single– cell-type mode, treating all super-pixels as a single population and identifying spatially coordinated transcriptional variation among neighboring pixels in an unsupervised manner (**Supplementary Fig. 4a-d**). The first CoPro component exhibited a strong and coherent spatial pattern across the tissue, with low and high scores localized near vascular structures and intermediate scores spanning the intervening parenchyma (**Fig. 6c, Supplementary Fig. 4e**). Examination of canonical zonation markers confirmed that this component faithfully recapitulated the known periportal-to-pericentral metabolic axis: periportal genes were enriched at one end of the gradient, while pericentral markers increased monotonically toward the opposite end (**Fig. 6d**; **Supplementary Fig. 4f-h, Supplementary Table 9**). These results demonstrate that CoPro can robustly recover the dominant lobular zonation axis using only imputed hepatocyte expression and super-pixel location, despite the absence of discrete cell boundaries and the presence of substantial technical noise.

Having established that CoPro reliably recovers hepatocyte zonation in this setting, we next asked whether it could uncover coordinated spatial programs between hepatocytes and endothelial cells, which are known to play a central role in establishing and maintaining zonation. We identified super-pixels that could be confidently assigned to hepatocytes or endothelial cells based on established marker genes, and applied CoPro in a cross–cell-type setting to identify spatially aligned transcriptional programs between these two populations.

Using CoPro’s kernel-weighted regression framework (**Methods**), we were able to transfer zonation scores from hepatocytes to endothelial cells (**Fig. 6e**). To interpret the endothelial contribution to this shared spatial program, we identified endothelial genes whose expression varied systematically along the inferred zonation axis. This analysis recovered well-established zonated endothelial morphogens, including the pericentral Wnt ligands *Wnt2, Wnt9b,* and *Rspo3* (**Fig. 6f, Supplementary Table 10**), consistent with their known role in reinforcing hepatocyte β-catenin signaling and pericentral identity^32,34^.

To independently validate these findings and assess whether the recovered endothelial zonation patterns reflect biological signal rather than imputation artifacts, we compared our results to a prior paired-cell RNA sequencing (pcRNA-seq) study^35^, in which endothelial gene expression was mapped to zonal identity through physical pairing with hepatocytes. Endothelial genes identified by CoPro as zonation-associated showed strong concordance with pcRNA-seq–derived zonation patterns (**Fig. 6g**, **Supplementary Fig. 5a-b**), including key morphogens and signaling factors. This agreement provides external validation that CoPro accurately recovers coordinated endothelial–hepatocyte spatial programs, even when applied to histology-imputed spatial transcriptomic data.

Together, these analyses establish that CoPro robustly recovers both the canonical hepatocyte metabolic zonation axis and its coordinated endothelial signaling programs in an exceptionally challenging data modality. This validated lobular axis provides a reliable spatial reference for subsequent analyses, enabling us to ask how aging-associated transcriptional programs are organized relative to baseline liver architecture.

### Aging liver: reproducible spatial aging modules and expansion of the midlobular zone

Having demonstrated that CoPro can be applied to pixel-level histology-imputed transcriptomic data to recover coordinated transcriptional programs along the liver lobular axis, we next asked how aging-associated transcriptional programs emerge on top of this baseline tissue architecture. We analyzed liver tissue from *Ercc1*^−/Δ^ mutant mice, a well-established model of accelerated aging characterized by genomic instability, cellular senescence, inflammation, and extracellular matrix remodeling^36^. Recent studies of liver aging suggest that aging does not manifest as a uniform perturbation across hepatocytes; instead, multiple aging-associated programs emerge within spatially structured niches across the lobule. For example, recent spatial and single-cell analyses have shown zonation-dependent metabolic changes and localized senescent niches associated with immune and fibroblast interactions during liver aging^37,38^. These observations raise the possibility that multiple spatially organized aging programs coexist within the same tissue. Such spatial heterogeneity makes CoPro particularly well-suited for decomposing overlapping transcriptional programs in aging liver.

We applied CoPro to six histology-imputed spatial transcriptomic datasets from *Ercc1*^−/Δ^ mutant livers (three female and three male). For each sample, we computed the top eight components to capture independent axes of coordinated spatial variation (**Fig. 7a**, **Supplementary Fig. 7-12**). To interpret these components biologically, we performed pathway enrichment analysis on the gene weights associated with each component. Pathways were grouped into biologically meaningful themes including senescence/SASP signaling, immune and interferon responses, proliferation/cell-cycle programs, extracellular matrix (ECM) remodeling and fibrosis, and stress-related pathways (**Supplementary Table 11**). Components were then compared across samples by computing similarities between their pathway enrichment profiles. Hierarchical clustering based on Euclidean distances of normalized enrichment scores grouped the components into transcriptional modules (**Supplementary Fig. 13**). Importantly, components originating from different animals frequently clustered together, indicating that these modules represent reproducible biological programs rather than sample-specific artifacts. Further details of the computational pipeline and enrichment analysis are provided in the **Methods**.

**Figure 7.**
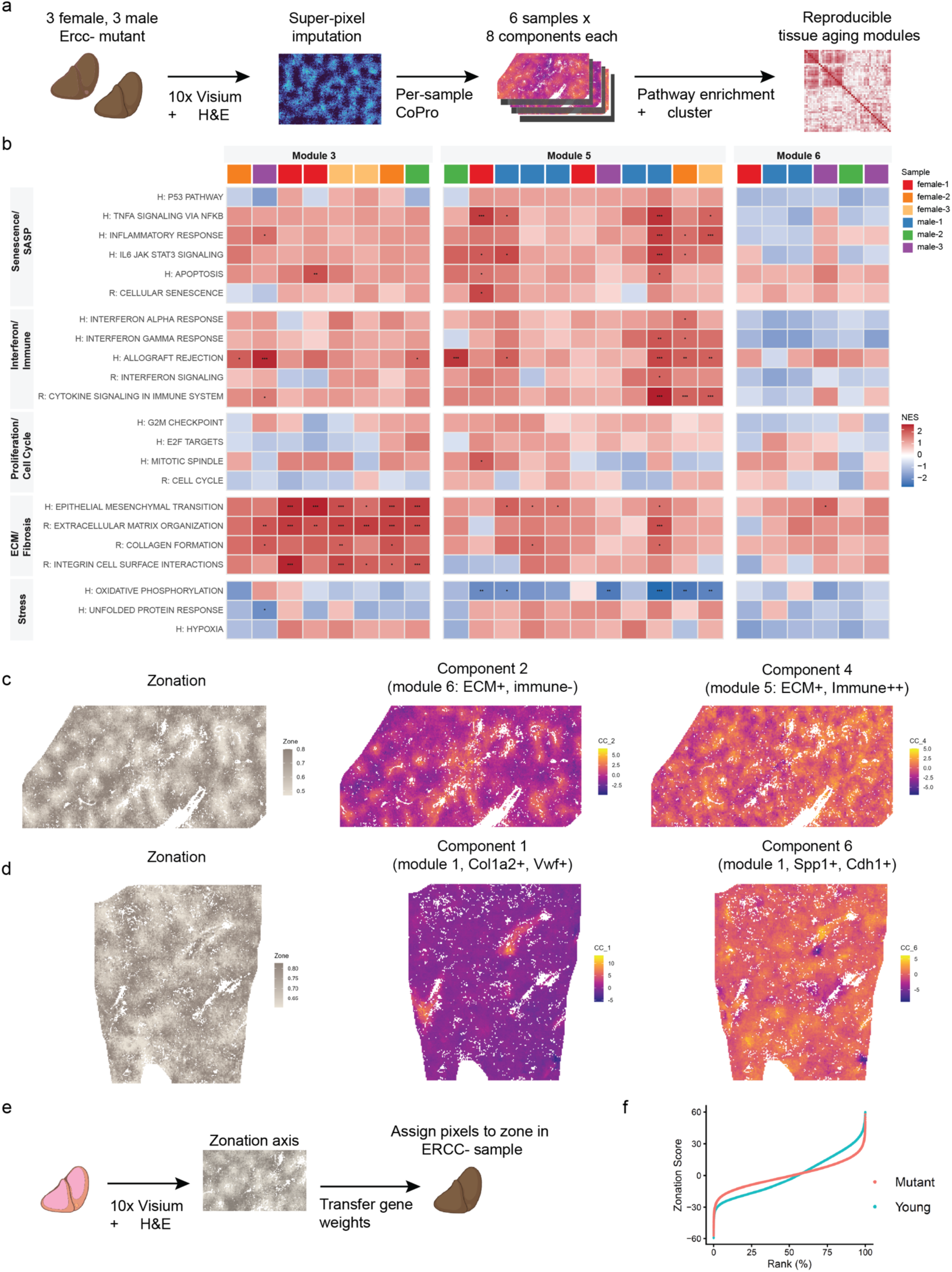
Reproducible spatial aging modules and expansion of the midlobular zone in mouse model of liver aging. **(a)** Overview of the analysis pipeline. Spatial transcriptomic data from six *Ercc1*^−/Δ^ mutant mouse livers (three female and three male) were obtained using 10x Genomics Visium and processed with superpixel imputation to increase spatial resolution. CoPro was applied independently to each sample to extract eight spatially coordinated transcriptional components. Gene weights for each component were interpreted using pathway enrichment analysis, and components were compared across samples by clustering their pathway enrichment profiles to identify reproducible transcriptional modules. **(b)** Three modules that are consistently observed across animals identified through cross-sample clustering of component pathway enrichment profiles. Heatmap shows normalized enrichment scores (NES) for pathway signatures associated with each component. Stars indicate statistical significance of pathway enrichment (***=0.0001, **=0.001, *=0.01). (**c,d**) Example of distinct aging modules within a single liver (c, male replicate 3; d, female replicate 3). (**e**) Transfer of the canonical zonation axis from healthy to mutant liver. The gene weights defining the primary CoPro zonation axis learned from healthy liver were fixed and applied to the mutant samples to compute zonation scores for each superpixel. This procedure uses the healthy liver as a spatial reference and enables direct comparison of zonation structure between healthy and aged tissues. (**f**) Zonation score by rank for each cell. Mean midlobular marker expression indicates expansion of zone 2 in mutant mouse liver compared to wild-type reference.

**Figure 7b** highlights three modules that were particularly prominent across samples: Modules 1, 4, and 5. The expanded component-by-pathway heatmap, including all 48 components, reveals additional smaller clusters that represent variations of these dominant programs (**Supplementary Fig. 14**). The most striking cluster corresponds to Module 1, which is characterized by extremely strong enrichment for pathways associated with extracellular matrix remodeling and fibrosis, together with moderate enrichment for interferon/immune and senescence/SASP signaling. This module shows large positive enrichment scores for multiple ECM-related pathways, including epithelial–mesenchymal transition, extracellular matrix organization, collagen formation, and integrin signaling, often accompanied by highly significant p-values. A second major cluster, Module 5, also exhibits ECM pathway enrichment but with stronger activation of immune and inflammatory signaling compared to Module 1, including interferon responses and cytokine signaling pathways. This pattern suggests a transcriptional program reflecting immune-driven tissue remodeling, in which matrix remodeling occurs alongside inflammatory activation. A third module, Module 4, also shows enrichment for ECM-related pathways but with little or no enrichment for immune or interferon signaling, indicating a structural ECM remodeling program with minimal immune involvement. These modules, therefore, distinguish multiple ECM-associated aging programs that differ in the degree of inflammatory activation.

We next examined how these modules relate to the lobular zonation axis, which represents the primary spatial gradient in liver physiology. For each component, we quantified the relationship between component scores and zonation scores using local spline smoothing. The resulting spline fits are shown in **Supplementary Figs. 15-17**. While individual components often displayed zonal preferences, such as periportal enrichment, pericentral enrichment, or non-monotonic patterns, these patterns were not consistent across samples within a given module. Components belonging to the same module could appear in different zones across animals. This indicates that the modules represent shared biological programs whose spatial realization varies between livers, rather than programs strictly tied to a fixed lobular zone. This observation is consistent with recent reports that aging-related hepatocyte states and senescent niches can appear in different lobular contexts depending on physiological or pathological conditions^37–39^.

To illustrate how CoPro dissects spatially overlapping aging programs within individual livers, we highlight two representative samples in **Fig. 7c–d**. **Figure 7c** shows two components from male replicate 3 together with the zonation reference map. The zonation panel provides the anatomical context of the lobule, enabling spatial interpretation of the aging programs. In the spatial maps, warm colors indicate high component scores and cool colors indicate low scores; spatially coherent color gradients reflect structured transcriptional programs. Component 2 belongs to Module 6, the ECM-dominant program with minimal immune activation. Spatially, this component is localized primarily near the pericentral region (**Supplementary Figs. 15-17**). In contrast, Component 4 belongs to Module 5, the immune-associated ECM-remodeling module, which in this sample localizes to both pericentral and periportal zones. Although Component 4 overlaps partially with the ECM-rich regions identified by Component 2, it extends more broadly across the tissue. These two components, therefore, reveal distinct but related tissue states, separating structural ECM remodeling from immune-driven matrix remodeling within the same liver.

Another example of within-sample heterogeneity is illustrated in female replicate 3 (**Fig. 7d**), where two components belonging to the same ECM-dominated module nevertheless capture distinct transcriptional programs (**Supplementary Fig. 18**). Component 1 is driven by classical extracellular matrix and fibrosis genes, including *Col1a1, Col1a2, Col6a3, Bgn, Sparc,* and *Timp2*, together with growth factor and vascular regulators such as *Tgfb2, Pdgfa, Edn1,* and *Vegfc*, indicating a program associated with structural matrix deposition and stromal remodeling. Spatially, this component is highly localized, forming discrete hotspots concentrated near pericentral regions. In contrast, Component 6 is dominated by a different set of genes that become prominent when the component weights are sign-aligned (i.e., considering genes with negative CC6 weights). These include *Spp1, Igfbp3, Igfbp7, Plaur, Serpine2, Dram1,* and *Aatk*, which are associated with senescence signaling, stress responses, and ECM regulatory pathways rather than structural collagen deposition. Unlike the focal pattern of Component 1, this program is spatially broader and more diffuse across the tissue. The two components separate structural ECM accumulation from a senescence-associated signaling program that regulates matrix remodeling, despite sharing enrichment for ECM-related pathways. This example illustrates how CoPro can resolve multiple overlapping transcriptional states within the same tissue, distinguishing ECM deposition from senescence-associated ECM remodeling.

In addition to identifying aging-associated transcriptional programs, we asked whether aging alters the integrity of the underlying lobular zonation itself. To directly quantify changes in liver spatial organization, we transferred the canonical zonation axis learned from healthy liver to the *Ercc1*^−/Δ^ mutant samples, analogous to our transfer of the crypt–villus axis from healthy to diseased colon (**Fig. 7e**). This approach uses the healthy liver as a spatial reference and enables direct comparison of zonation structure across conditions.

When canonical zone-specific marker genes were plotted along the transferred zonation axis, we observed systematic changes in zone composition and boundary sharpness in aged liver (**Fig. 7f**). Markers of the midlobular zone (zone 2) exhibited a marked expansion along the axis, extending toward both the periportal (zone 1) and pericentral (zone 3) ends. In contrast, zone 1 and zone 3 markers occupied narrower ranges of the axis, indicating a relative contraction of these terminal zones. Moreover, the transitions between zone 1 and zone 2, and between zone 2 and zone 3, were substantially less abrupt than in healthy liver, with overlapping expression of zone markers across broader intervals of the axis. Together, these results indicate that aging leads to expansion of the midlobular zone and progressive blurring of zonal boundaries.

Taken together, these analyses reveal that liver aging involves two concurrent but distinct processes: the emergence of multiple spatially structured aging-associated transcriptional programs and a progressive loss of precision in canonical lobular zonation. By using the healthy liver as a spatial reference, CoPro distinguishes changes to the underlying tissue scaffold from aging programs that are superimposed upon it, enabling an interpretable decomposition of tissue-level changes during aging.

### CoPro reveals corticomedullary vascular specialization in the kidney

The mammalian kidney is organized along the corticomedullary axis, where nephron epithelium follow a stereotyped anatomical progression from proximal tubule (PTS1–3) in the cortex through the loop of Henle and thick ascending limb to the distal convoluted tubule and connecting tubule (DCT-CNT) back in the cortex^40,41^ (**Fig. 8a**). This tubular architecture is organized alongside a complex vascular network spanning glomerular capillaries (GC), efferent and afferent arterioles (EA, AA), cortical peritubular capillaries (PTC), and the descending and ascending vasa recta (DVR, AVR) of the medulla^42,43^. Prior work established that each vascular compartment expresses distinct transcription factors, solute transporters, and angiocrine factors — including zone-restricted transporters such as *Slc14a1* and *Aqp1* in the DVR and *Slco4a1* in the AVR — but required extensive developmental profiling, antibody staining, and lineage-tracing experiments across multiple studies to define this zonation hierarchy^44,45^. Whether the full transcriptional continuum of vascular specialization could be recovered directly from spatial transcriptomic data, without prior knowledge of vascular subtype markers, remained untested.

**Figure 8.**
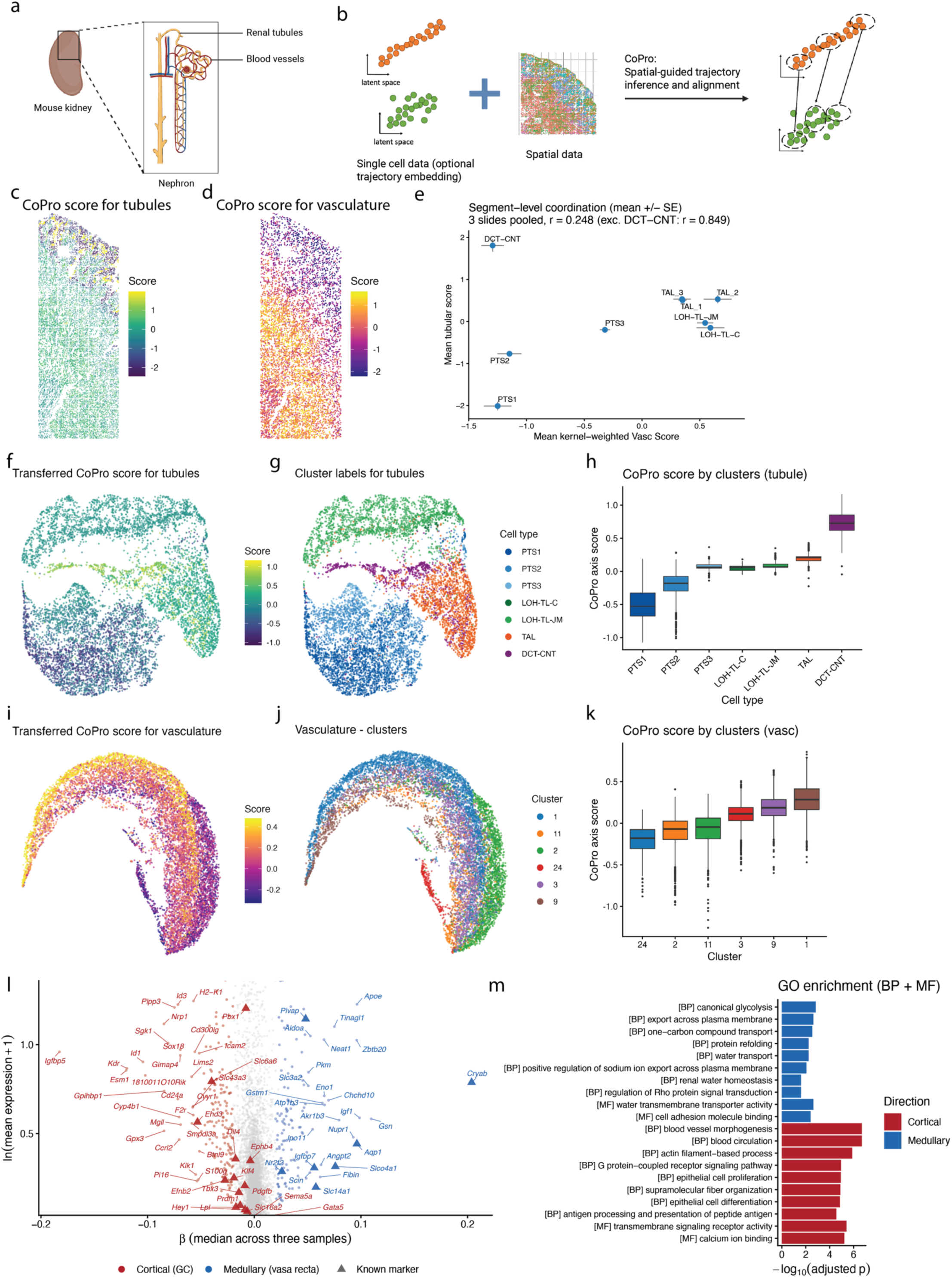
CoPro reveals corticomedullary vascular specialization in the kidney using supervised spatial co-progression. **(a)** Schematic of kidney tissue organization. Nephron segments follow a stereotyped anatomical progression along the corticomedullary axis, accompanied by a parallel vascular network spanning glomerular capillaries to the vasa recta. **(b)** Overview of the supervised CoPro workflow. The known anatomical ordering of tubular segment types is encoded as an ordinal reference axis from single-cell data. CoPro integrates this reference with spatial data to identify the vascular gene program maximally co-progressing with nephron position through the spatial kernel. **(c–d)** Spatial projection of CoPro scores for tubular cells **(c)** and vascular cells **(d)** on a representative kidney tissue section. Tubular scores recapitulate the corticomedullary gradient from proximal tubule (low) to collecting duct (high). Vascular scores reveal a parallel spatial gradient distinguishing cortical from medullary endothelial populations. (**e**) Segment-level coordination between tubular and vascular axes. Each point represents the mean kernel-weighted vascular score versus the mean tubular score for a nephron or vascular segment, pooled across three biological replicates (mean +/- SE). The overall correlation is r = 0.248; excluding DCT-CNT, which anatomically loops back toward the cortex, the correlation increases to r = 0.849. (**f-g**) UMAP of tubular epithelial cells colored by transferred CoPro score (**f**) and annotated tubular cell type (**g**). (**h**) Box plots of transferred CoPro scores stratified by tubular cell type, confirming monotonic ordering from PTS1 through TAL segments, with DCT-CNT scoring high on the tubular axis. (**i-j**) UMAP of vascular cells colored by transferred CoPro score (**i**) and vascular cluster label (**j**). (**k**) Box plots of transferred scores by vascular cluster, revealing a graded separation across six endothelial populations. (**l**) MA plot of full-transcriptome regression identifying genes associated with the vascular corticomedullary axis. Median regression coefficients (β) across three biological replicates are plotted against mean expression (ln(mean expression + 1)). Genes concordant across all three replicates (same sign, FDR < 0.05 in ≥ 2) are colored by direction: cortical/glomerular-enriched (red, negative β) and medullary/vasa recta-enriched (blue, positive β). Known vascular zone markers from Barry et al. are shown as triangles and labeled; genes with |β| > 0.05 are also labeled. (**m**) Gene Ontology enrichment analysis (Biological Process and Molecular Function) of concordant axis-associated genes.

To address this, we applied CoPro in supervised mode to kidney seqFISH data comprising 1,298 genes across three biological replicates^46^. The known anatomical ordering of 11 tubular segment types was encoded as an ordinal reference axis, which CoPro used to identify the vascular gene program maximally co-progressing with nephron position through the spatial kernel (**Fig. 8b**, **Supplementary Table 12**). The supervised tubular axis recovered the known nephron ordering with good agreement (Kendall tau = 0.74–0.78 across replicates). Visualization of cell scores in situ confirmed that tubular scores recapitulate the corticomedullary gradient, with proximal tubule cells scoring lowest and collecting duct cells scoring highest (**Fig. 8c**). The coordinating vascular scores revealed a parallel spatial gradient, with cortical endothelial cells (Vasc_1) scoring distinctly from medullary endothelial cells (Vasc_2), demonstrating that CoPro captured corticomedullary vascular specialization directly from spatial co-expression patterns (**Fig. 8d, Supplementary Table 13**). We next characterized the segment-level concordance between the tubular and the kernel-weighted vascular scores. With the notable exception of DCT-CNT cells, the tubular and vascular cells show a strong positive correlation (r = 0.85 excluding DCT-CNT): despite carrying the highest tubular scores, DCT-CNT cells are surrounded by low-scoring cortical vasculature because they anatomically loop back to the cortex after the loop of Henle’s descent into the medulla (**Fig. 8e**). This decoupling may also partly reflect the limited spatial resolution of 2D tissue sections, where the fine-scale boundaries between PT- and DCT-adjacent vasculature are difficult to resolve without 3D profiling. Results were highly concordant across replicates (pairwise Pearson r = 0.77–0.92 for tubular gene weights, 0.62–0.74 for vascular gene weights).

To extend vascular axis discovery beyond the spatial gene panel, we transferred CoPro’s vascular gene weights to a kidney scRNA-seq atlas^47^ (31,265 cells; **Methods**). Transferred scores organized nephron epithelial cells (n = 9,117) along the expected corticomedullary ordering in UMAP space, and similarly resolved vascular cells (n = 10,198) into a continuous cortical-to-medullary gradient spanning six endothelial clusters, although these clusters have not been mapped to specific anatomic structure in the original study (**Fig. 8f–k**). Full-transcriptome regression against the transferred scores identified 3,485 zonation-associated genes concordant across all three replicates (same sign and FDR < 0.05 in at least two replicates) (top genes shown in **Fig. 8l**). Strikingly, the recovered axis recapitulated the complete vascular zonation hierarchy established by prior developmental studies, despite being derived entirely from spatial co-expression with tubular cells rather than from endothelial subtype annotations.

Examining regression coefficients for genes with known vascular zone specificity revealed great concordance (23 of 24 testable genes, 96%) with previously mapped expression patterns (**Supplementary Fig. 19**). At the cortical/glomerular end, GC transcription factors (*Gata5*, *Tbx3*, *Prdm1*), efferent arteriole transporters, angiocrine factors, and renin-angiotensin system components were enriched. At the medullary end, the axis captured venous and vasa recta identity, including *Nr2f2*, all three vasa recta transporters (*Slc14a1*, *Aqp1*, *Slco4a1*), and medullary angiocrine factors. The single discordant gene (*Ephb4*) had a small effect size near zero, suggesting a borderline rather than contradictory assignment (**Supplementary Fig. 19**).

Gene ontology analysis of concordant axis genes revealed functionally coherent programs at each pole (**Fig. 8m**). Genes enriched at the cortical/GC end were associated with blood vessel morphogenesis, blood circulation, epithelial cell proliferation, and actin filament-based processes, consistent with the active angiogenic and vascular remodeling programs of glomerular capillaries and cortical peritubular vasculature. Genes at the medullary/vasa recta end were enriched for glycolysis, transmembrane transport, water homeostasis, and protein refolding, reflecting both the specialized metabolic and transport functions of the vasa recta and the cellular stress responses characteristic of the hyperosmotic, hypoxic medullary environment.

Together, these results demonstrate that CoPro’s supervised mode, guided solely by the known nephron anatomy and the principle that neighboring cell types coordinate their transcriptional programs, infers the hierarchy of renal vascular specialization from spatial transcriptomic measurement — with limited gene panels and without endothelial subtype annotations. The approach provides a general framework for using established tissue architecture as a biological prior to discover coordinated gene programs in neighboring cell types.

## Discussion

We present CoPro, a computational framework for detecting spatially coordinated progression of cellular states across cell types in spatial transcriptomic data. Several distinguishing features make CoPro well-suited to questions that existing tools cannot readily address. First, CoPro operates at single-cell resolution and does not require cells of different types to be co-aggregated into shared spatial bins, which prevents data loss for rare or spatially sparse cell populations. Second, the spatial kernel-restricted CCA objective explicitly models intercellular coupling driven by spatial proximity, enabling CoPro to identify interpretable multi-dimensional tissue gradients. Importantly, CoPro adaptively selects the spatial kernel bandwidth from the data, allowing cell– cell interactions to be captured at the appropriate spatial scale without manual tuning. Third, the model and estimation procedure effectively separate multiple orthogonal axes of coordinated variation that overlap in space, as demonstrated by our decomposition of crypt morphology and acute injury responses in Day 3 colon, and of distinct aging-associated transcriptional programs in aged liver. Fourth, the transfer of gene weights from a reference condition to additional samples defines a common biological axis across samples without relying on spatial registration, making cross-sample comparison possible even in tissues with high morphological variability.

Applying CoPro to four tissue types demonstrated these capabilities across diverse biological settings. In the mouse colon undergoing DSS-induced injury, CoPro recovered the epithelial crypt differentiation axis at homeostasis and an orthogonal acute early stress response for samples taken at early disease. Transferring the homeostatic crypt-villus gene weights to later disease timepoints allowed us to quantify the progressive disruption of intercellular spatial alignment during peak inflammation and its differential recovery at Day 21. Mapping samples taken during injury to a common CoPro axis allowed the quantification of disease severity across mice, validated by manual annotation.

In the liver, CoPro robustly recovered hepatocyte lobular zonation from histology-imputed transcriptomic data at 8-μm resolution—a setting where conventional clustering and trajectory methods failed—and identified coordinated endothelial zonation programs validated by orthogonal paired-cell RNA-seq data. In aged mutant liver, CoPro decomposed three biologically distinct aging-associated programs superimposed on the canonical zonation axis and revealed that aging produces both the emergence of spatially structured senescence states and a progressive blurring of zonal boundaries.

In the brain, CoPro enabled the dissection of overlapping spatial gradients that are difficult to resolve using existing approaches. In the striatum, where both D1 and D2 medium spiny neurons exhibit continuous transcriptional variation, standard dimensionality reduction and spatial methods tend to recover a single dominant gradient. By leveraging flexible kernel design, CoPro decomposed this variation into distinct dorsal–ventral and medial–lateral axes, each associated with coherent and biologically interpretable gene programs. These results demonstrate that CoPro can resolve multiple, spatially overlapping axes of organization within the same cell populations, providing a more nuanced view of brain tissue architecture than approaches that collapse variation into a single manifold.

In the kidney, CoPro’s supervised mode enabled the inference of coordinated transcriptional programs across anatomically linked but transcriptionally distinct cell types. Using the known ordering of nephron segments as a reference axis, CoPro identified a corresponding vascular program that co-progresses with epithelial position along the corticomedullary axis. This analysis recovered the full continuum of vascular specialization from cortical to medullary endothelium without requiring prior annotation of vascular subtypes. Notably, the inferred vascular axis recapitulated known molecular markers established through extensive experimental studies, demonstrating that CoPro can leverage spatial coupling to infer coordinated molecular transitions across cell types.

Analysis of the colon injury dataset also illustrates the key design choices underlying CoPro’s multi-sample integration strategy. For biological or technical replicates with limited sample-to-sample variation, multiple samples can be jointly analyzed to identify shared axes of spatially-coordinated progression across cell types, as demonstrated with the three Day 0 replicate samples. When substantial sample-to-sample variation is expected, e.g. differing disease severity across mice, CoPro can instead be applied to a representative sample, and the estimated gene weights can be transferred to the remaining samples to define a common interpretable axis. This shared axis allows direct comparison of cellular co-progression across samples, as demonstrated through our decomposition of disease progression scores across the Day 9 samples.

We acknowledge that the transfer-based integration strategy has inherent limitations. By projecting all samples onto a common axis derived from a single reference, the approach may obscure tissue-specific patterns that exist in individual samples but deviate from the reference program. The transfer strategy anchors integration to biologically meaningful gene weights rather than raw expression values, but at the cost of imposing a common reference frame on all samples. More broadly, CoPro inherits limitations common to linear factor models: the CCA objective captures linear co-variation, so strongly nonlinear co-progression patterns may require future extensions. Additionally, the quality of the recovered axes depends on the accuracy of input cell type annotations, and computational cost scales with the number of cell pairs in the kernel, which may require subsampling strategies for very large datasets.

The CoPro framework is not inherently limited to transcriptomic features. The spatial kernel-restricted CCA objective operates on any low-dimensional cell-level representation, and could in principle be applied to features extracted from other data modalities that are defined at cellular or spatial resolution. One particularly promising direction is the direct application of CoPro to morphological features derived from H&E-stained tissue images. Computational pathology methods increasingly produce high-quality, spatially resolved cellular embeddings from routine histology slides, and CoPro could leverage these embeddings to detect coordinated morphological progression across cell types without requiring any transcriptomic measurement. This would open the framework to the vastly larger body of existing H&E data and could serve as a complement or precursor to spatial transcriptomic profiling. Extending CoPro to such modalities, and to settings combining morphological and transcriptomic features in a single analysis, represents a natural avenue of interest for future work.

## Methods

### Input data

Spatial transcriptomic data were represented as a normalized gene-expression matrix and spatial coordinates for each annotated cell type. For cell type *t*, we denote expression by *Z*_*t*_ ∈ *R*^*n*_*t*_×*m*^ and spatial coordinates by *L*_*t*_. Expression was centered and variance-scaled within cell type, followed by principal component analysis (PCA):

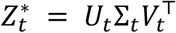

The corresponding PC score matrix was used as the low-dimensional representation; in the default CoPro pipeline, PCs were further variance-normalized (whitened) before optimization.

### Distance-based spatial Kernel

CoPro incorporates the spatial information by constructing a spatial kernel that reflects pairwise cell interactions. The vanilla spatial kernel is constructed through a Gaussian kernel function applied to Euclidean distance matrix (*d*_*E*_) with a bandwidth parameter, *σ*.

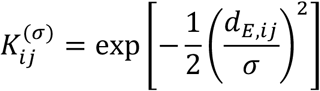

For tissues with complex topology, an alternative kernel is a morphology-aware kernel that integrates Euclidean proximity and graph geodesic structure. Briefly, we computed Euclidean distances between cells and geodesic distances on a *k*-nearest-neighbor graph (default k = 10) built in spatial coordinates. A linear model relating Euclidean to geodesic distance was fit over local neighborhoods, and cell pairs that were Euclidean-close but geodesically distant were reclassified as effectively non-local by assigning a large distance. This yielded a filtered distance matrix *d*_*f*_ that we can use in place of *d*_*E*_.

### Spatial kernel restricted CCA (skrCCA)

Given *T* cell types, CoPro estimated canonical directions {*w*_*t*_}^*T*^ by maximizing kernel-weighted cross-type covariance in PC space:

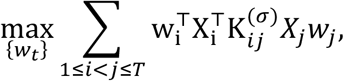

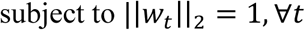

where *X*_*t*_ is the cell-type-specific PC embedding. CoPro provides two optimization strategies for solving this objective. The default is block-coordinate power iteration: for each cell type *i*, the weight vector is updated as *w*_*i*_ ← normalize(∑_j❜*i*_*X*_*i*_^*T*^*K*_*i*j_*X*_j_*w*_j_), cycling through all cell types until convergence. An alternative coordinate descent strategy is also available, in which each update is a convex combination of the current weight and the power iteration direction: *w*_*i*_ ← normalize((1 − *α*)*w*_*i*_ + *α* ⋅ normalize(*update*)), where *α* ∈ (0, 1) is a step-size parameter. The coordinate descent approach provides more stable convergence in settings where the standard power iteration may oscillate, such as when additional constraints or regularization are applied. For within-cell-type analysis (single cell type), the objective reduces to maximizing the quadratic form *w*^*T*^(*X*^*T*^*KX*)*w* subject to unit-norm constraints, where K is the within-cell-type spatial kernel; the optimum is the leading eigenvector of this matrix without iteration. Additional canonical components were obtained by sequential deflation, yielding orthogonal higher-order axes of coordinated variation.

### Multi-sample modeling framework

For multi-sample analyses, CoPro treats each sample as a separate observation of the same underlying cross-cell-type coupling structure. Let *q* ∈ {1, …, *Q*}index samples and *t* ∈ {1, …, *T*} index cell types. For each sample *q* and cell type *t*, CoPro computes a sample-specific PC embedding *X*_*t*,*q*_while maintaining a common feature basis across samples within each cell type. For each cell-type pair (*i*, *j*), sample-specific spatial kernels 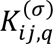 are constructed at bandwidth *σ*.

Rather than estimating sample-specific canonical directions, CoPro estimates shared canonical weight vectors 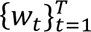 across all slides by maximizing the summed cross-slide objective:

objective:

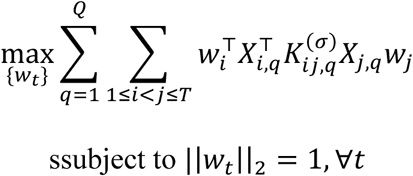

This pooling strategy improves robustness and promotes the recovery of biological axes that are reproducible across tissue sections, rather than patterns idiosyncratic to individual samples. In practice, CoPro alternates block-coordinate updates of each *w*_*t*_: each update aggregates contributions from all sample-level kernel-weighted cross-covariance terms involving that cell type, followed by 𝑃_2_-normalization. Convergence is declared when the maximum change in any weight vector falls below a user-specified tolerance.

Higher-order components are obtained via sequential deflation: the variance explained by previously identified components is removed, and the shared-weight objective is re-optimized on the residual. Each canonical axis is thus defined globally at the dataset level, while per-sample cell scores are obtained by projecting onto the sample-specific embeddings:

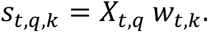

Downstream summaries—such as normalized or bidirectional spatial correlations—can then be reported either per sample or as aggregate statistics across samples.

### Kernel bandwidth selection

We evaluated skrCCA over a grid of kernel bandwidths *σ*. For each cell-type pair (*i*, *j*) and component *k*, we computed a kernel-normalized canonical correlation:

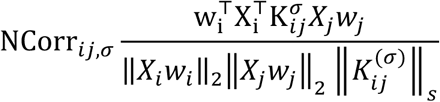

Where ǁ*K*_*i*j_ǁ_*s*_ denotes the spectral norm of the kernel matrix. The optimal bandwidth was defined as the *σ* that maximizes the mean first-component normalized correlation across cell-type pairs.

We note that although NCorr defined here has nice statistical properties (bounded from [-1, 1] and order-invariant), it is not a correlation measure per se, hindering an intuitive interpretation of the strength of cellular coordination in terms of the actual correlation. Thus, we introduced a Bidirectional Spatial Correlation, defined as

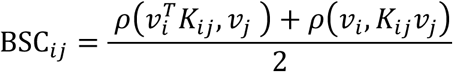

where *ρ* stands for the Pearson Correlation Coefficient.

### Identification of coordinated gene programs

Because PCA and skrCCA are linear, canonical weights were projected back to gene space to obtain gene loadings for each component and cell type. Cell-level canonical scores were computed as *s*_*t*,*k*_ = *X*_*t*_*w*_*t*,*k*_, interpreted as continuous co-progression coordinates CC_k. In addition to PCA back-projection, CoPro provides an alternative regression-based approach for obtaining gene-level weights: for each gene, expression is regressed against the cell score, and the resulting coefficient is used as the gene weight. This approach avoids collinearity issues inherent in PCA back-projection and yields more reproducible gene weights across biological replicates. To assess gene associations with these latent states while controlling for covariates and sample-level heterogeneity, we used linear mixed-effects regression in a DIALOGUE-like framework as an optional downstream analysis^10^.

### Identification of co-progressed cell states

The canonical variable *X*_*t*_*w*_*t*_can be viewed as a new feature space that represents a continuous cell state, which we call “cell score”. For a model with multiple orthogonal canonical variables detected, we refer to each cell score as CC_k, *k in* {1, 2, …, *K*}.

### Permutation-based significance testing

To assess statistical significance of each canonical axis, CoPro implements a spatial permutation procedure. The tissue is partitioned into non-overlapping spatial bins (default side length equal to twice the kernel bandwidth σ). Bins are then randomly shuffled, and for each cell type being tested, the cells at each bin’s locations are replaced by same-type cells sampled from the mapped target bin, while spatial coordinates remain fixed. This preserves cell-type identity and coarse spatial structure at the bin scale while disrupting the fine-grained spatial coupling between cell types that CoPro detects. The kernel-restricted CCA is re-estimated on the permuted data to build a null distribution, and p-values are computed as the fraction of permutation replicates exceeding the observed normalized correlation.

### Projection of identified axes to another slide

To project an axis identified in a reference dataset onto a target sample, CoPro transfers gene-space loadings and computes projected cell scores within each cell type. Let 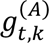 denote the reference gene loadings for cell type *t* and component *k*, and 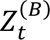 denote the target expression matrix. Target expression is transformed using reference-derived feature statistics: (i) optional per-gene quantile normalization of target to the reference distribution; and (ii) centering and scaling by the reference mean and standard deviation. Projected cell scores are then computed as

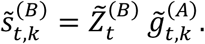

For multi-cell-type analyses, projection is performed independently per cell type and subsequently aggregated at the object level.

Projection fidelity is assessed in the target sample using target-slide kernels evaluated at the chosen *σ*: (i) kernel-normalized cross-type correlation, computed as described above with *X*_*t*_*w*_*t*,*k*_replaced by 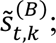 and (ii) bidirectional spatial correlation metrics capturing both cross-type and within-type (self-kernel) spatial structure. For multi-sample targets, these metrics are reported either per sample or as aggregate summaries across samples.

### Simulation of spatial transcriptomics data with or without coordinated progression

We used scRNA-seq data from the Liver Cell Atlas^23^ to create the simulated spatial transcriptomics dataset. We first generate random spatial positions for cells in a two-dimensional tissue region and assign each cell a random score, based on a standard normal, and a random type label, based on a multinomial distribution with predefined probabilities. We then performed spatial smoothing to induce a spatial gradient, which we use as the ground truth cell scores. Of note, the spatial smoothing can be conducted either within individual cell types or across cell types. When spatial smoothing is limited to cells within the same type, spatial autocorrelation is introduced within that cell type, but not co-progression between cell types. This serves as our null hypothesis. Spatial smoothing of cell scores across types introduces spatial co-progression, giving us the alternative hypothesis. To assign realistic expression profiles to these positions, we compute PCA on the normalized and scaled expression data and generate random weight vectors on the principal components. These weights define the ground truth gene program associated with the spatial gradient. Ground truth cell scores were then calculated for each cell in the single-cell dataset based on gene weights and matched with the scores on the simulated spatial data, creating a cell-by-gene expression matrix paired with spatial coordinates. This approach ensures that the simulated data retains realistic expression distributions and gene-gene correlations while embedded into a known spatial gradient.

### Simulation of orthogonal spatial co-progression axes

For these simulations, we generated two independent smooth spatial gradients by simulating two random initial scores per cell position, applying spatial smoothing with a Gaussian kernel separately to each, and then explicitly orthogonalized the second axis by regressing out the first. We used optimal transport-based matching on quantile-transformed cell scores to assign cells to spatial positions. Since two-dimensional matching is more complicated and may distort the original target spatial pattern (despite the high correlation itself), we conducted a PC regression on the cell score to assign the estimated cell score as the ground truth for this analysis.

### Comparison with other methods GASTON

Spatial gradients were inferred using GASTON, a neural network–based manifold learning framework that jointly models gene expression and spatial coordinates. For each simulated dataset, gene expression matrices (Parquet format) and matched spatial metadata were loaded into a Scanpy AnnData object. Expression values were library-size normalized, log-transformed, and reduced by principal component analysis (PCA), retaining 50 components. The PCA embedding (proxy for GLM-PCA) and 2D spatial coordinates were exported as tab-delimited inputs to Gaston.

Models were trained using a two-branch architecture with spatial hidden layers [64, 32] and expression hidden layers [128, 64]. Training was performed for 3000 epochs with checkpoints every 500 epochs and one random seed per replicate. Each simulation replicate was processed independently using an automated batch framework to ensure consistent parameterization across scenarios.

### SpaceFlow

SpaceFlow is a graph-based deep learning framework that integrates gene expression similarity with spatial proximity. Expression matrices and spatial metadata were loaded into an AnnData object, with genes detected in fewer than three cells removed. SpaceFlow preprocessing retained the top 3000 highly variable genes.

Models were trained with spatial regularization strength 0.1, latent dimension 50, learning rate 1×10⁻³, and 1000 training epochs, using early stopping (patience = 50; minimum epochs = 100). An edge subset size of 1,000,000 was used for graph regularization. All runs were performed in CPU mode with a fixed random seed (42). The learned low-dimensional embedding was exported automatically. Spatial domains were identified using nearest-neighbor graph clustering (n_neighbors = 50, resolution = 0.5), and pseudo-spatiotemporal maps were computed using a graph-based approach (n_neighbors = 20, resolution = 0.5).

### ONTraC

Gene expression matrices were library-size normalized, log-transformed, and reduced to principal components computed from highly variable genes prior to graph construction. ONTraC constructed a k-nearest neighbor graph (k = 6) in reduced-dimensional space and trained a graph neural network to model directional cellular progression. The model was optimized for up to 1,000 epochs with early stopping (patience = 100) using a learning rate of 0.03 and batch size of 5. The objective function integrated modularity, regularization, and purity constraints (modularity weight = 0.3, regularization weight = 0.1, purity weight = 300) with a smoothness parameter (β = 0.03) to promote coherent lineage structure. Cells were assigned continuous pseudotime values and discrete lineage branches inferred from graph topology.

### Mouse brain data analysis with CoPro

We applied CoPro to MERFISH data from the mouse brain^48^, focusing on D1 and D2 medium spiny neurons in the striatum. To decompose overlapping spatial gradients along distinct anatomical axes, we leveraged CoPro’s flexible kernel design by constructing axis-specific spatial kernels. For the dorsal–ventral axis, we used a Gaussian kernel that weighted cell pairs preferentially along the dorsal–ventral direction while down-weighting lateral separation, and conversely for the medial–lateral axis. This directional kernel design enabled independent recovery of orthogonal gene expression gradients that standard isotropic kernels would conflate into a single dominant axis.

### Mouse colon data analysis with CoPro

#### Dataset and preprocessing

We downloaded the processed mouse colon dataset from Data Dryad. In this study, we focused on the interactions within the mucosa layer, so we retained the epithelial, fibroblast, and macrophages within this layer.

#### Mapping all Day 9 samples to a unified axis

Day 9 samples displayed great structural and expression-level variabilities, reflecting different severity of diseases. Here, we hypothesized that each sample capture different stages of a unifying disease/inflammatory axis. This inspired us to identify the disease axis within a slide displaying mild infection (ID: *100221_D9_m5_1_slice_2*) and transfer the gene weights to other slides. To retain comparability, we also self-transferred the gene weights to the reference slides so all cell scores are on the same scale.

#### Per-slide score aggregation and correlation with MU8 neighborhood abundance

To validate that the unified disease progression axis captures meaningful variation in disease severity across tissue sections, we correlated per-section CoPro scores with the abundance of the MU8 neighborhood—the most severely inflamed mucosal microenvironment identified in the original study.

For each of the 25 Day 9 tissue sections, transferred disease progression scores were aggregated across all three cell types (epithelial, fibroblast, and immune) by taking the 90th percentile per-cell score across all cells in the section. MU8 abundance per section was quantified as the fraction of all mucosal cells assigned to the MU8 neighborhood label in the original annotation, computed as the number of MU8-labeled cells divided by the total number of cells across all eight mucosal neighborhoods (MU1–MU8). Pearson correlation between the per-section 90th percentile disease score and the per-section MU8 fraction was then computed across all 25 sections (**Fig. 5g**). To further characterize within-section spatial variation, each tissue section was additionally partitioned into overlapping local spatial windows (side length 100 spatial units, 50% overlap in both dimensions). For each window, the 90th percentile disease score (aggregated across all three cell types) and the proportion of each cell type were computed.

#### Partitioning variance in disease progression scores

To quantify sources of variation in disease progression scores across the Day 9 cohort, we fit an intercept-only linear mixed-effects model using the **lme4** package in R, with mouse identity as a top-level random effect and tissue section nested within mouse as a second-level random effect:

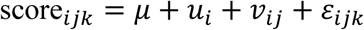

where 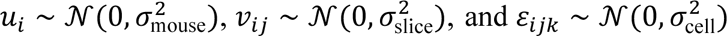 denote mouse-, section-, and cell-level residuals, respectively. Variance components were extracted via VarCorr(), and the proportion of total variance attributable to each level was computed as 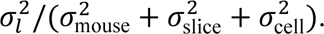 Under this parameterization, 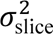 captures residual between-section variation within a mouse after accounting for the mouse-level mean; the relative magnitude of mouse versus section effects is therefore an empirical finding rather than a structural property of the model. The finding that mouse-level variation exceeds section-level variation indicates that systemic, animal-wide differences in disease severity—rather than spatial sampling of distinct tissue regions within the same animal—are the primary driver of inter-sample heterogeneity.

#### Identifying cell-type-specific genes associated with disease severity

To identify cell-type-specific genes associated with disease severity, we fit a separate linear mixed-effects model for each gene–cell-type combination using **lmerTest**:

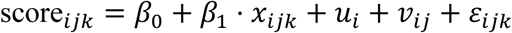

where *x*_*i*j*k*_ is the normalized expression of a given gene and the structure of random effects matches that above. The coefficient *β̂*_1_ and its *p*-value, derived via Satterthwaite’s degrees-of-freedom approximation, were recorded for each gene. *P*-values were corrected for multiple testing using the Benjamini–Hochberg false discovery rate (FDR) procedure, and significant genes were defined by FDR <0.001, mean expression > 0.1, and ∣ *β̂*_1_ ∣> 0.5.

#### Assessing the within- and cross-type coordination of cell programs across time

To quantify the spatial coordination of cell programs along the homeostatic crypt–villus axis across disease time points, we first transferred gene weights learned from Day 0 samples to all remaining samples (Days 3, 9, and 21). For each cell type, transfer proceeded in two steps: (i) optional feature-wise quantile normalization of the target sample to match the reference (Day 0) expression distribution; (ii) standardization of the quantile-normalized target using per-gene means and standard deviations estimated from the reference. Per-cell scores in the target sample were then obtained by projecting the standardized expression matrix onto the filtered weight matrix.

We next computed the Bidirectional Spatial Correlation (BSC) as a measure of how tightly cells co-progress along the transferred axis in space. Cells with negligible kernel support (row or column sums below 5 × 10^-2^) were excluded prior to computation to prevent instability from near-isolated cells. BSC values—both self- and cross-type—were computed per tissue section for each of the four time points and all cell type pairs. For multi-section comparisons, values were reported both per section and as within-mouse medians to account for the hierarchical sample structure.

### Mouse liver data analysis with CoPro

#### Dataset and preprocessing

We analyzed paired H&E-Visium data from wild-type and *Ercc1*^-/Δ^ mutant mice liver. To overcome the limited 55-μm spatial resolution of Visium technology, we applied the iSTAR method to computationally impute gene expression at 8-μm resolution from H&E images. This imputation process creates analytical challenges: each imputed pixel may contain partial or mixed cellular information, and H&E staining variability can introduce artifacts.

To ensure data quality, we applied stringent filtering: gene expression values were capped at the 95th percentile to reduce outlier effects, and spots with log total counts beyond mean ± 3 standard deviations were removed to eliminate technical artifacts. Following quality control, we normalized expression values using logCP10K normalization (log(counts per 10,000 + 1)), a standard approach for sparse transcriptomic data.

#### Cell type annotation and filtering

We used established marker genes to identify pixels corresponding to specific cell types. For hepatocyte analysis (representing ∼70% of liver cells), we retained pixels with high expression of hepatocyte markers. For endothelial cell analysis, we selected pixels with high endothelial marker expression and low hepatocyte marker expression to minimize contamination from adjacent hepatocytes in the imputed data.

#### Recovery of liver zonation patterns using within-cell-type CoPro

We divided one slide into training and test halves to evaluate the robustness of CoPro. Given the large number of hepatocyte pixels in the training half, we further partitioned it into two pseudo-slides to enable computational tractability while preserving spatial relationships (**Supplementary Fig. 4a**). We applied the multi-sample version of CoPro to these two pseudo-slides, using the within-cell-type mode to identify coordinated transcriptional programs within hepatocytes. This analysis generated cell scores capturing the dominant spatial transcriptional gradient. We then validated the recovered zonation pattern by: (1) examining spatial correspondence between cell scores and known lobular architecture (central veins and portal triads), (2) correlating cell scores with canonical zonation markers, and (3) transferring learned patterns to the held-out test half to assess generalization.

#### Visualization of zonation marker expression trends

To visualize how canonical zonation markers vary along the CoPro-inferred spatial axis, we plotted smoothed expression trends for representative periportal, midlobular, and pericentral gene sets. Super-pixels with CoPro scores outside the mean ± 2 standard deviations were excluded as outliers. Expression values were then min-max normalized to [0, 1] as (*x* − min (*x*))/(max (*x*) − min (*x*)). Zone-group summary curves were computed by averaging normalized expression across all genes assigned to each zone (periportal, midlobular, or pericentral). Smooth trends were fitted using a generalized additive model (GAM) as a function of the CoPro zonation score and displayed without 95% confidence intervals. The same procedure was applied to individual genes to generate per-gene trend plots for supplementary figures.

#### Comparison with paired-cell RNA sequencing data

To validate that the endothelial zonation patterns recovered by CoPro reflect biological signal rather than imputation artifacts, we compared our results against published paired-cell RNA sequencing (pcRNA-seq) data^35^ (GSE84498), in which endothelial gene expression was mapped to lobular zone identity through physical co-localization with adjacent hepatocytes. For each gene in the pcRNA-seq dataset, we computed a center-of-mass (COM) score from the per-zone mean expression:

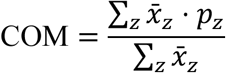

where *x̄*_*z*_ is the mean expression in zone *z*and *p*_*z*_ = (*z* − 1)/(*Z* − 1) maps zone indices linearly to [0, 1] from pericentral to periportal. COM scores thus capture each gene’s abundance-weighted position along the lobular axis. Expression was subsequently normalized to each gene’s maximum zonal mean. The top 20 spatially zonated endothelial genes identified by CoPro (ranked by gene weight magnitude) were selected for comparison. A back-to-back heatmap was then constructed with genes ordered by their spatial-data-derived COM scores: the left panel shows LOESS-smoothed expression across the CoPro-inferred zonation score from the super-pixel data, and the right panel shows pcRNA-seq normalized zonal mean expression for the same genes in the same order, enabling direct visual assessment of concordance between the two independent datasets.

#### Projecting endothelial cells onto the hepatocyte zonation gradient

To examine endothelial molecular signatures across liver zones, we projected endothelial spots onto the hepatocyte-defined zonation axis using CoPro’s cross-cell-type kernel framework. Specifically, we computed endothelial cell scores by multiplying hepatocyte cell scores with the hepatocyte-endothelial cross-type kernel matrix, effectively transferring spatial positional information from hepatocytes to endothelial cells based on their spatial proximity.

To identify zonation-associated genes in endothelial cells while minimizing contamination from adjacent hepatocytes in the imputed data, we focused on endothelial-specific genes. Following the approach from paired-cell RNA sequencing (pcRNA-seq) studies, we defined endothelial-specific genes as those with ≥2-fold higher expression in endothelial cells compared to hepatocytes in a reference single-cell dataset. We then fit quadratic regression models between each endothelial-specific gene’s expression and the projected zonation score to identify genes whose expression varied systematically across the periportal-pericentral axis.

### Decomposition of aging-associated spatial programs

We analyzed Visium spatial transcriptomics data from six *Ercc1*^−/Δ^ mutant mouse liver samples (three female, three male; approximately 4–4.5 months of age). For each sample, histology-imputed pixel-level gene expression counts and spatial coordinates were extracted. Pixels flagged by tissue masking were removed, and the remaining tissue area was spatially subsetted to a contiguous rectangle of approximately 40,000 pixels using an adaptive aspect-ratio search that minimized empty space within the selected region. Pixels below the 5th percentile of total UMI counts were removed, and genes with zero variance across pixels were excluded. Pixels with total counts exceeding 10 median absolute deviations from the median were removed as outliers. No library-size normalization or log-transformation was applied; raw counts were used as input, as CoPro performs internal centering and scaling during its PCA step. After filtering, each sample retained approximately 21,000–37,000 pixels and ∼998 genes.

CoPro was run independently on each sample using the single-sample interface. Principal component analysis was performed on the centered and scaled expression matrix, retaining the top 40 components. Pairwise Euclidean distances between pixel spatial coordinates were computed, and Gaussian kernel matrices were constructed at three bandwidth values (σ = 0.02, 0.05, and 0.1), with an upper quantile cutoff of 0.85. Sparse kernel CCA was then applied with a maximum of 500 iterations to extract eight canonical components per sample. Normalized correlations were computed for each component as a quality metric. Cell scores (per-pixel component loadings) and gene scores (per-gene component weights) were extracted at σ = 0.05 for downstream analysis.

For each component, Pearson correlations were computed between cell scores and three reference signals: a zonation score derived from known pericentral, midlobular, and periportal marker genes; a senescence signature based on the mean expression of curated senescence-associated genes; and a fibroblast signature based on curated fibroblast marker genes.

Gene scores for each component were used as ranking statistics for gene set enrichment analysis using fgsea (fgseaMultilevel, 10,000 permutations, gene set size 5–500). Enrichment was assessed against 33 curated pathway gene sets from MSigDB (Hallmark and Reactome collections), organized into seven biological themes: Senescence/SASP, Interferon/Immune, Proliferation/Cell Cycle, ECM/Fibrosis, Angiogenesis/Endothelial, Stress/Oxidative/UPR, and Metabolic/Zonation. Two additional custom gene sets (senescence and fibroblast marker genes) were also tested. P-values were adjusted using the Benjamini–Hochberg procedure.

To identify reproducible spatial expression programs across samples, we constructed a normalized enrichment score (NES) profile for each of the 48 components (8 per sample × 6 samples) across the 33 curated pathways. Hierarchical clustering was performed on the Euclidean distance matrix of NES profiles using Ward’s method (ward.D2), and the dendrogram was cut to yield 10 modules. Within each module, a sign-alignment procedure was applied: the initial consensus NES profile was computed as the column-wise mean across module members, and any component whose NES profile was negatively correlated with the consensus had its NES values and cell scores multiplied by −1. This procedure was iterated until convergence (up to three iterations) to ensure coherent orientation of components within each module. Module zonation profiles were visualized by plotting sign-aligned cell scores against the zonation coordinate. Consensus pathway enrichment profiles were summarized as the mean NES across module members, with standard errors and one-sample t-test p-values reported for each pathway.

### Mouse kidney data analysis with CoPro

#### Kidney seqFISH data processing

We analyzed kidney seqFISH data from three healthy control mouse kidney tissue sections^46^ (Ctrl1, Ctrl2, Ctrl3). The dataset profiled 1,298 genes across 107,161 total cells with 22 annotated cell types. For CoPro analysis, we retained tubular epithelial cells (PTS1, PTS2, PTS3, LOH-TL-C, LOH-TL-JM, TAL_1, TAL_2, TAL_3, DCT-CNT, PC, IC) and vascular endothelial cells (Vasc_1, Vasc_2, Vasc_3), comprising 63,740 tubular and 18,185 vascular cells across the three samples. Log-normalized gene expression values from the Seurat^49^ Spatial assay were used as input.

#### Supervised CoPro analysis

To define the reference tubular axis, we assigned ordinal labels reflecting the known anatomical ordering of nephron segments: PTS1 = 1, PTS2 = 2, PTS3 = 3, LOH-TL-C = 4, LOH-TL-JM = 4, TAL_1–3 = 5, DCT-CNT = 6, PC = 7, IC = 7. LOH-TL-C and LOH-TL-JM were assigned the same label because they represent cortical and juxtamedullary segments of the same thin limb structure at equivalent anatomical depth; similarly, PC and IC were grouped because both principal and intercalated cells reside within the collecting duct at the same tubular position. PCA was computed on tubular cells (40 components, centered and scaled), and the ordinal labels were regressed onto the PCA scores by ordinary least squares. The resulting coefficient vector (excluding the intercept) was normalized to unit length to define the supervised tubular weight vector **W**_1_^tub^, which was passed to CoPro’s supervised mode via the transferred_weight_1 parameter, fixing the first canonical component for tubular cells while allowing the vascular component to be optimized.

To derive the corresponding vascular weight vector, we computed a Gaussian spatial kernel between vascular and tubular cells (bandwidth *σ* = 0.1mm). The kernel-weighted mean of tubular axis scores was computed for each vascular cell, and these smoothed scores were regressed onto the vascular PCA scores to obtain **W**_1_^vasc^. CoPro was then run with both weight vectors fixed as first components.

Axis recovery quality was evaluated for each sample by Kendall’s rank correlation (**τ**) between CoPro cell scores and the ordinal segment labels.

#### Cross-platform transfer to scRNA-seq

To extend vascular gene program discovery beyond the seqFISH panel, we transferred CoPro’s vascular gene weights to a kidney scRNA-seq atlas comprising 31,265 cells and 19,125 genes post-QC^47^. Transfer used regression-based gene weights, which showed higher cross-replicate reproducibility than PCA back-projection weights. The 1,250 genes shared between the seqFISH panel and the scRNA-seq transcriptome were projected via CoPro’s transfer_scores function with quantile normalization, with the seqFISH vascular expression matrix serving as the reference distribution. Scores were transferred independently for each of the three control replicates and averaged to obtain a consensus score per cell.

#### Full-transcriptome regression

To identify axis-associated genes beyond the spatial panel, we regressed each gene in the full scRNA-seq transcriptome against the transferred axis scores by ordinary least squares, after excluding genes with standard deviation below 10^-5^ and mean-centering both expression values and axis scores. Regression coefficients (*β*) and significance were assessed via Pearson’s correlation test, with *P*-values corrected for multiple testing by the Benjamini-Hochberg procedure. Regression was performed independently for each control replicate. A gene was considered concordant if its *β* coefficient had the same sign across all three replicates and reached FDR < 0.05 in at least two of three.

#### Gene ontology enrichment analysis

Gene ontology (GO) enrichment analysis was performed on concordant axis-associated genes using the clusterProfiler R package (v4.18.4). Genes were stratified into positive-*β* (medullary end) and negative-*β* (cortical end) sets. Gene symbols were mapped to Entrez IDs using the org.Mm.eg.db annotation database, and over-representation analysis was performed against GO Biological Process and Molecular Function ontologies using enrichGO (Benjamini-Hochberg correction, *P*_adj_ < 0.05), with the full set of tested genes as the background. Redundant GO terms were collapsed using the simplify function (semantic similarity cutoff = 0.5).

## Data availability

We downloaded the following spatial transcriptomics datasets from public repositories:

Healthy and injured colon MERFISH data^6^: https://doi.org/10.5061/dryad.rjdfn2zh3

Brain MERFISH data^48^: https://knowledge.brain-map.org/data/5C0201JSVE04WY6DMVC/explore

Liver Cell Atlas data^23^: https://www.livercellatlas.org/

Liver pcRNA-seq data^35^: http://www.ncbi.nlm.nih.gov/geo/query/acc.cgi?acc=GSE84498

The aging liver Visium data were generated in-house and are part of a joint submission focusing on the underlying biology. These data will be publicly available upon publication of the companion paper or upon request to the corresponding authors.

## Code Availability

CoPro is available as open-source software in both R and Python. The R package can be installed from GitHub (https://github.com/Zhen-Miao/CoPro), and the Python package copro is available at https://github.com/Zhen-Miao/copro-python and on PyPI. Code for reproducing the analyses presented in this paper is also available at the GitHub repository.

## Supplementary Materials

**Supplementary Figures 1-19**

**Supplementary Table 1:** Conceptual comparison of methods for spatial transcriptomic analysis

**Supplementary Table 2-4:** Benchmark performance of CoPro and comparison methods in simulations (naïve, proportions, and orthogonal)

**Supplementary Table 5:** CoPro gene weights in healthy colon (Day 0). PCA back-projection and regression-based weights for Epithelial, Fibroblast, and Immune cells (CC1).

**Supplementary Table 6:** CoPro gene weights in DSS-treated colon (Day 3). PCA back-projection and regression-based weights for Epithelial, Fibroblast, and Immune cells (CC1 and CC2).

**Supplementary Table 7**: Cell-type-specific genes associated with disease severity in DSS-treated colon (Day 9)

**Supplementary Table 8:** CoPro gene weights in DSS-recovered colon (Day 21). PCA back-projection and regression-based weights for Epithelial, Fibroblast, and Immune cells (CC1).

**Supplementary Table 9:** CoPro gene weights in healthy mouse liver

**Supplementary Table 10:** Zonation-associated endothelial genes in healthy liver, quadratic

**Supplementary Table 11:** Pathway grouping for the mutant liver (*Ercc1*^⁻/Δ^)

**Supplementary Table 12:** CoPro gene weights in kidney tubular segments. PCA back-projection and regression-based weights.

**Supplementary Table 13:** CoPro gene weights in kidney vasculature. PCA back-projection and regression-based weights.

## Author Contribution

N.R.Z. and Z.M. conceived the study. Z.M. and N.R.Z. designed the statistical model. Z.M. implemented the model and constructed the software package with feedback from S.H., J.K., and N.R.Z. Y.Q. further improved the software for scalability. Z.M. and S.H. conducted the simulation data analysis. L.L. generated the liver Visium data. Z.Z. conducted the super-pixel imputation analysis for the liver data, and A.A. preprocessed the data. Z.M. and N.R.Z. conducted the real data analysis with guidance from L.L., J.R.M., and J.K.. N.R.Z. supervised the work. Z.M. and N.R.Z. wrote the manuscript with feedback from all coauthors.

## Acknowledgements

We thank Dr. Jiaoyang Huang for constructive discussion on the null distribution of the framework. We thank Dr. Jeffrey R. Moffitt and Dr. Paolo Cadinu for guidance on the analysis of the mouse colon injury data set. We thank all members in Nancy Zhang Lab and Junhyong Kim Lab for their constructive comments. We acknowledge funding from the NIH (R01GM25301, R01GM149671, and 1R56AG081351 to NRZ) and NSF (DMS/NIGMS grant DMS-2245575 to NRZ).

---

**Supplementary Figure 1.**
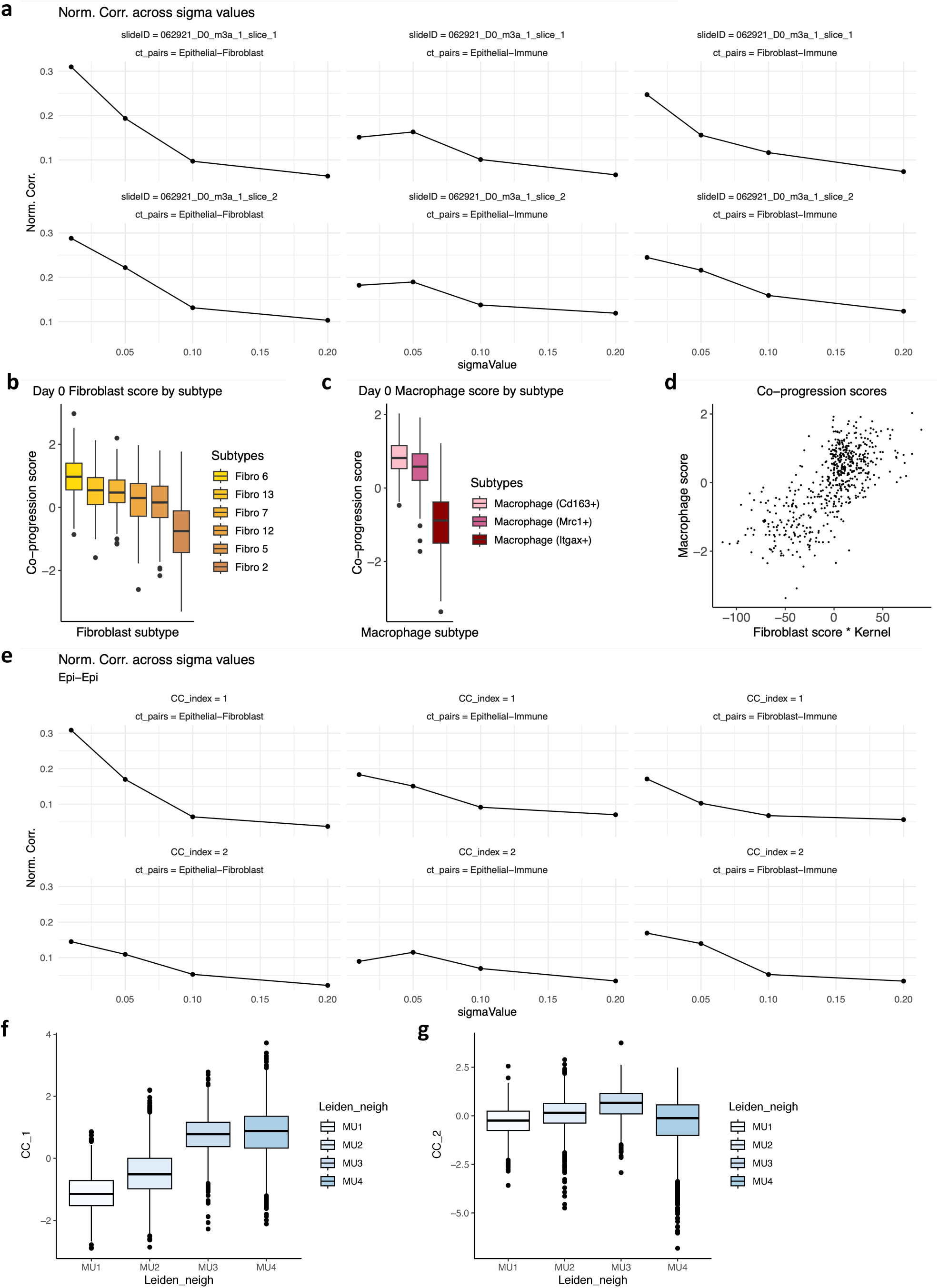
(**a**) Normalized correlation across sigma values for Day 0 colon co-progression analysis, evaluated for all three cell type pairs (Epithelial–Fibroblast, Epithelial–Immune, Fibroblast–Immune) across two representative tissue sections. (**b**) Fibroblast co-progression scores grouped by subtype at Day 0. The scoring links fibroblast 6 to the crypt base and fibroblast 2 to the crypt tip. (**c**) Macrophage co-progression scores by subtype at Day 0, revealing that Itgax+ macrophages localize toward the tip while Mrc1+ macrophages localize near the crypt base. (**d**) Scatter plot of kernel-smoothed co-progression scores between macrophage and fibroblast cell types in a representative Day 0 section (**e**) Normalized correlation across sigma values for Day 3 colon co-progression analysis, evaluated for all three cell type pairs (Epithelial–Fibroblast, Epithelial–Immune, Fibroblast–Immune) and two retained canonical components (CC1 and CC2). (**f–g**) Distribution of Day 3 CC1 (**f**) and CC2 (**g**) scores stratified by mucosal microenvironment (ME) neighborhood labels from the original publication.

**Supplementary Figure 2.**
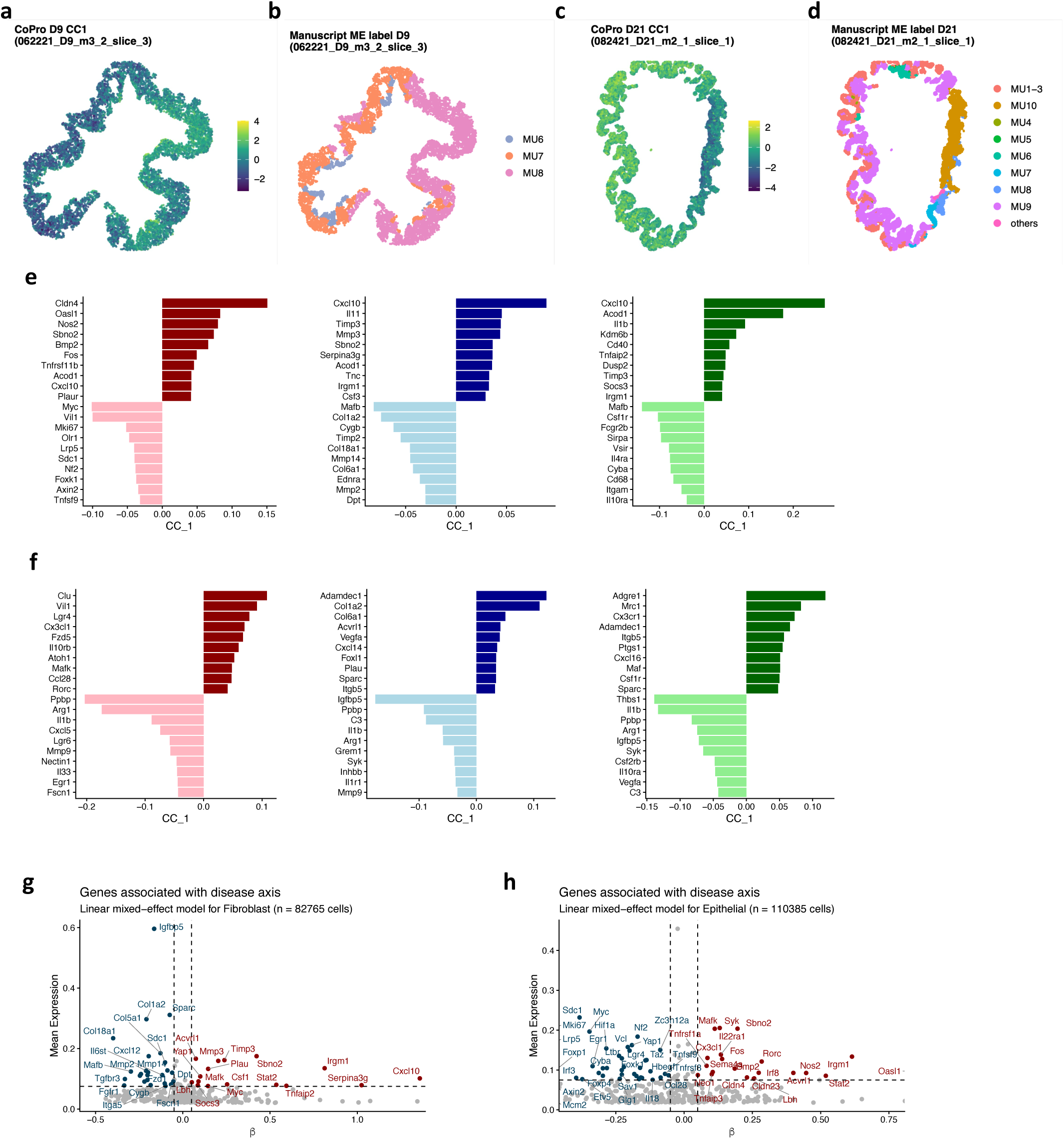
(**a-d**) Spatial maps projecting CoPro canonical component 1 (CC1) scores alongside the manuscript’s manually curated microenvironment (ME) labels for Day 9 (severe inflammation, **a-b**) and Day 21 (repair, **c-d**) samples. (**e-f**) Cell-type-specific gene weights derived from the CoPro progression axes for Epithelial, Fibroblast, and Immune cells at Day 9 (**e**) and Day 21 (**f**). (**g-h**) Identification of cell-type-specific genes associated with disease severity. Results map the linear mixed-effects model outputs for Fibroblast (**g**) and Epithelial (**h**) cells, adjusting for mouse- and slice-level variation.

**Supplementary Figure 3.**
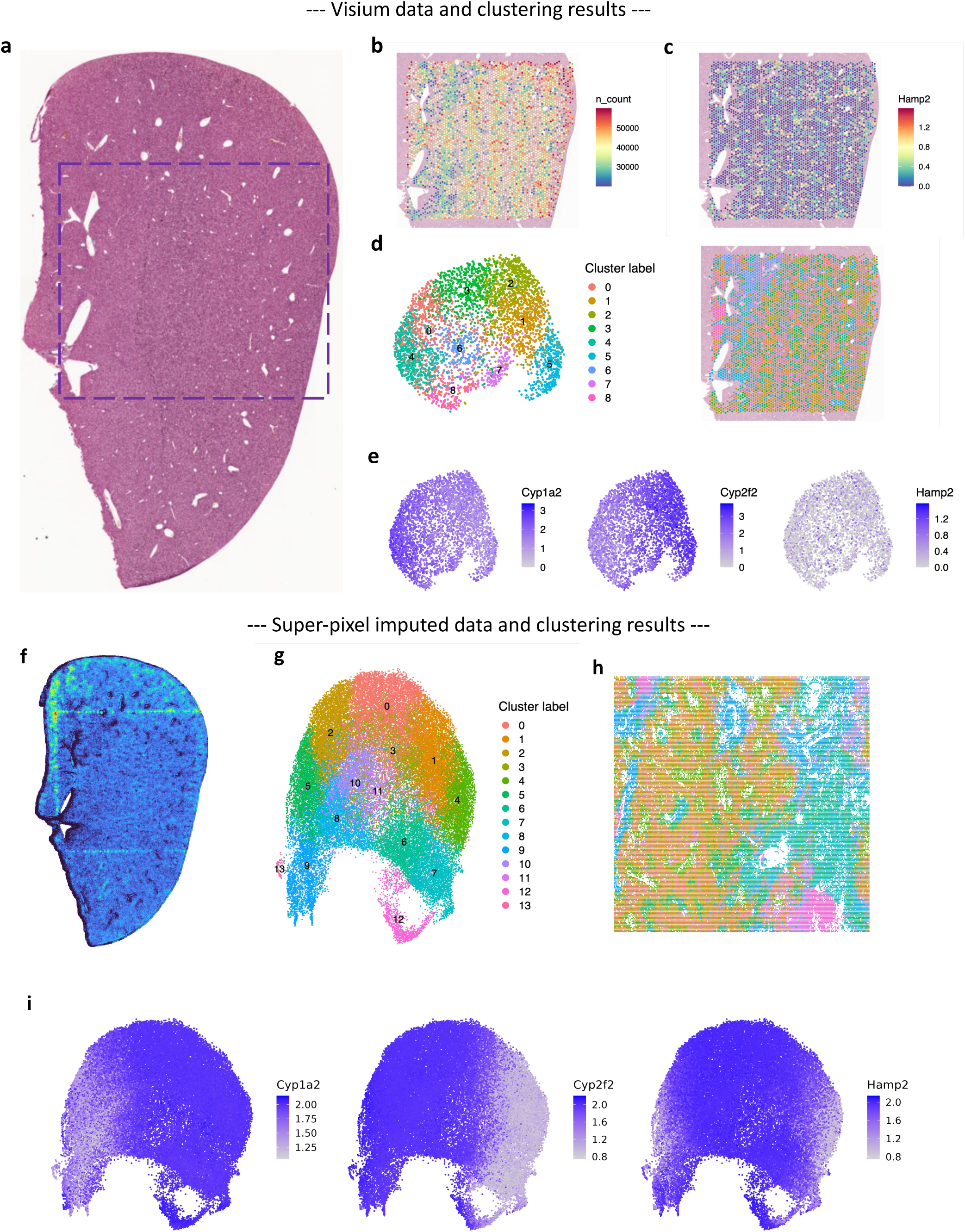
(**a**) H&E-stained image of a healthy wild-type mouse liver section. The purple rectangle marks the region captured by Visium experiment. (**b**) Spatial map of total UMI counts for Visium spots. (**c**) Spatial expression of a representative zonation marker (*Hamp2*) on Visium spots. (**d**) Leiden clustering of Visium spots projected onto tissue coordinates. (**e**) Spatial expression of canonical zonation markers on Visium spots. (**f**) *Hamp2* expression for the full liver section computed on 8-μm super-pixel imputed data (iSTAR) (**g-h**) Leiden clustering of iSTAR super-pixels, shown on UMAP (**g**) and tissue coordinates (**h**). (**i**) Spatial expression of canonical zonation markers on iSTAR super-pixels.

**Supplementary Figure 4.**
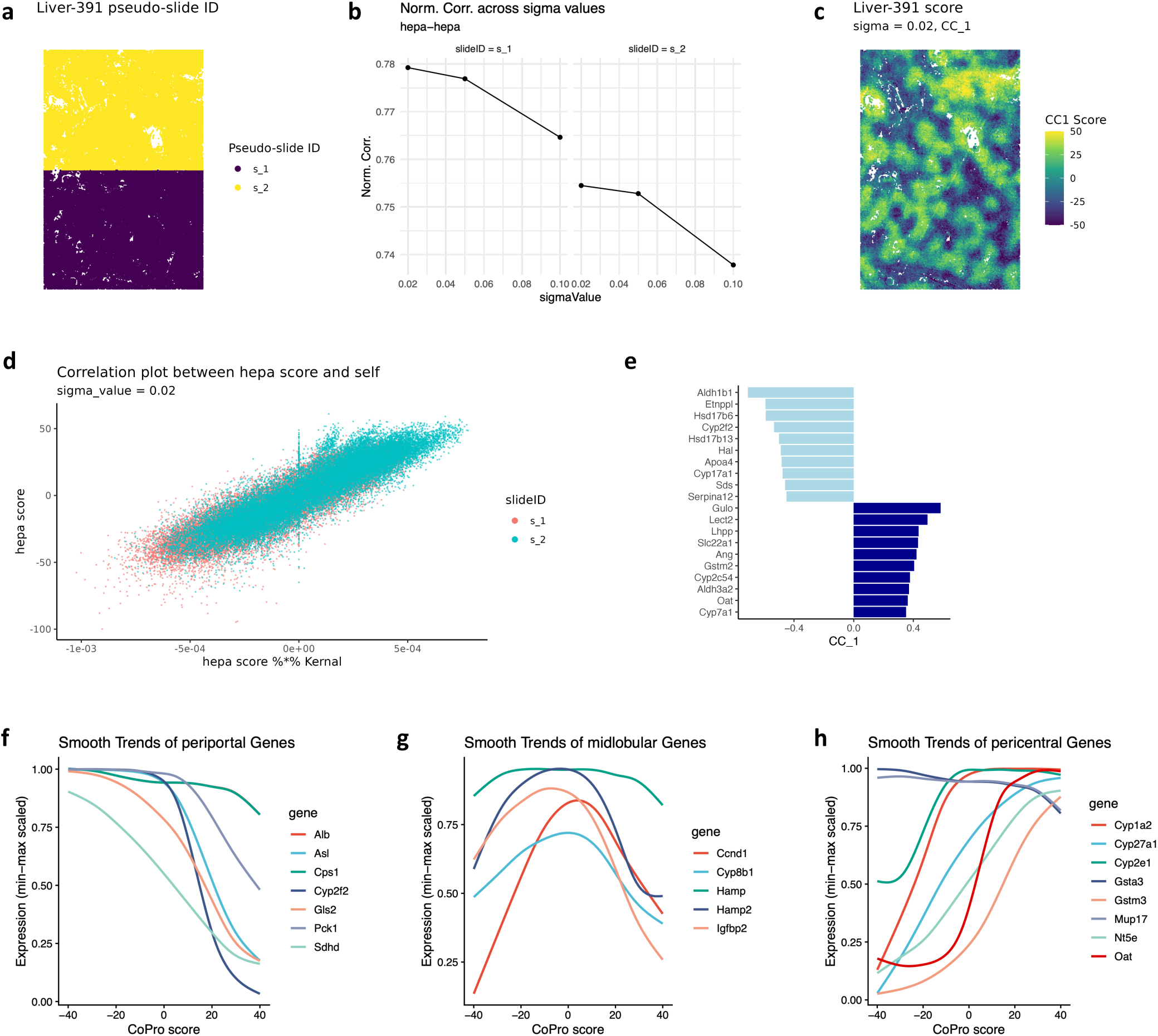
(**a**) Pseudo-slide partition of the Liver-391 training half that enables multi-slide CoPro analysis. (**b**) Normalized correlation across sigma values for single-cell-type hepatocyte CoPro analysis. (**c**) CoPro CC1 score map projected onto the Liver-391 section. The spatial pattern shows coherent periportal-to-pericentral organization, with high and low scores localized near vascular structures and intermediate scores spanning the intervening parenchyma. (**d**) Scatter plot of kernel-smoothed hepatocyte CC1 scores against the cell score, with each point colored by pseudo-slide identity (s_A vs s_B). (**e**) CC1 gene weight bar plot for hepatocytes. (**f-h**) Smooth expression trends of canonical zonation markers (f: periportal, g: midlobular, h: pericentral) as a function of CoPro score.

**Supplementary Figure 5.**
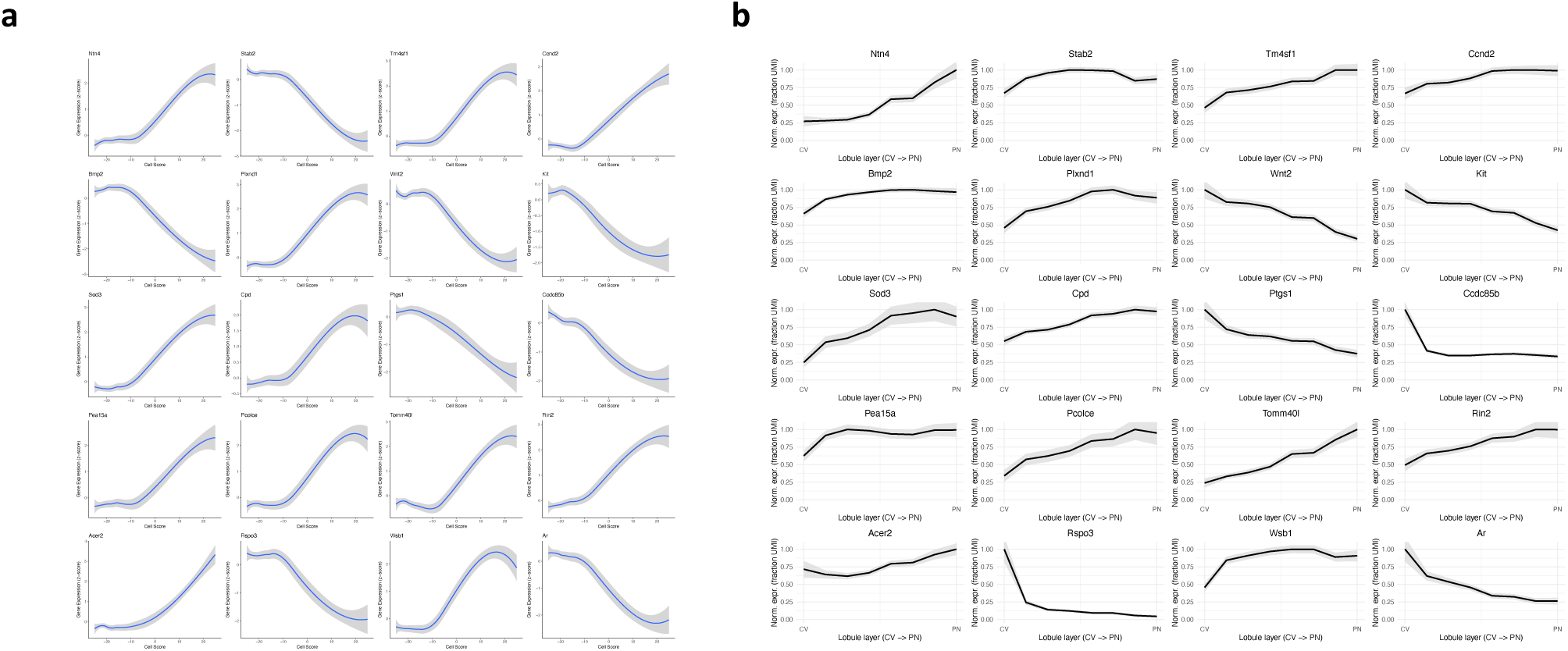
(**a**) Smooth trends of individual endothelial gene expression plotted along the CoPro-inferred zonation score axis. (**b**) Normalized zonal mean expression for the same genes from published paired-cell RNA sequencing (pcRNA-seq) data across lobular layers.

**Supplementary Figure 6.**
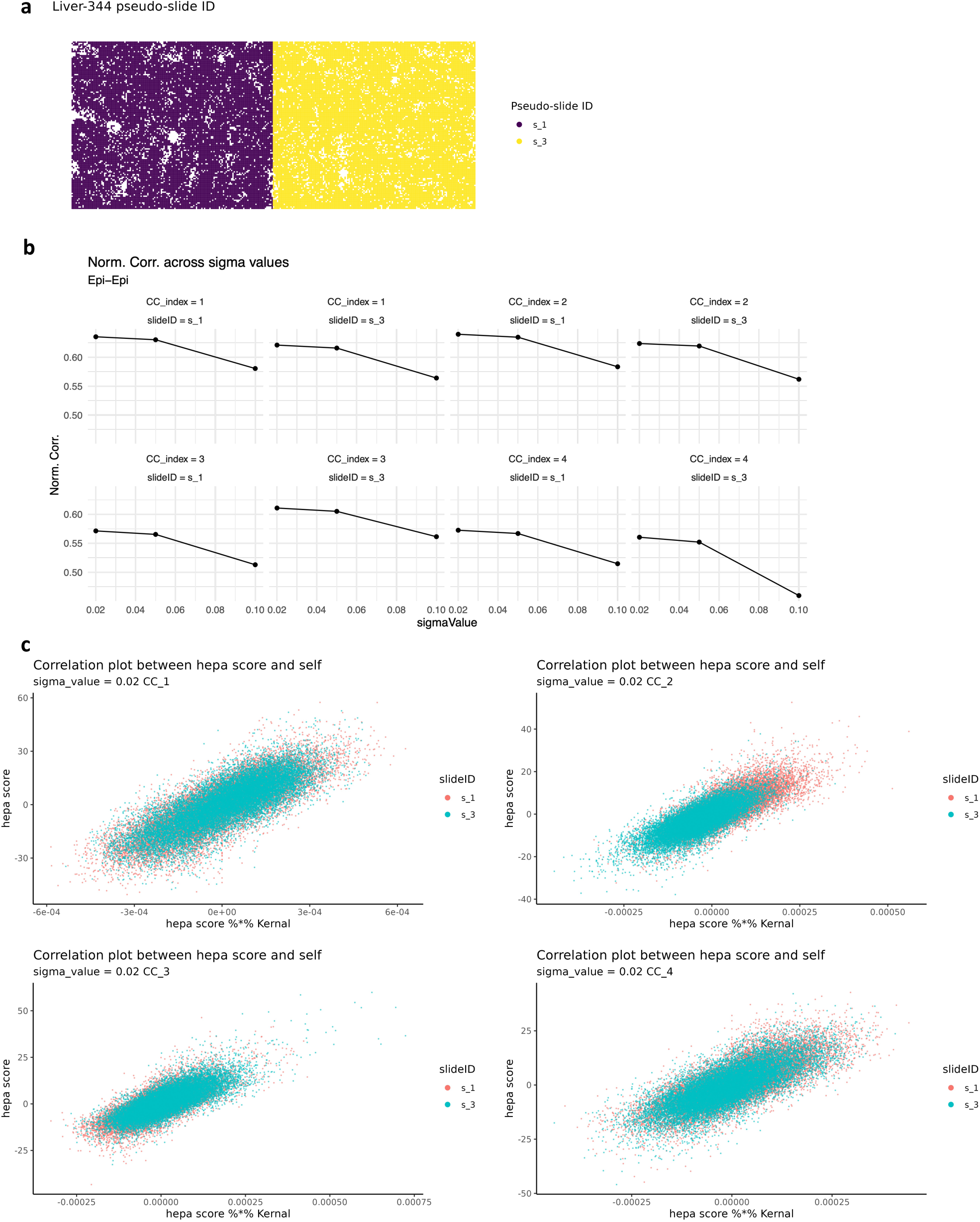
(**a**) Pseudo-slide partition of the Liver-344 data that enables multi-slide CoPro analysis. (**b**) Normalized correlation across sigma values for Liver-344 hepatocyte CoPro analysis, shown for all four canonical components (CC1–CC4) across all pseudo-slides. (**c**) Scatter plots of kernel-smoothed hepatocyte scores between pseudo-slide pairs for each of the four canonical components (CC1–CC4) at sigma = 0.02. In each panel, points are colored by pseudo-slide identity (s_1, s_3).

**Supplementary Figure 7-12.**
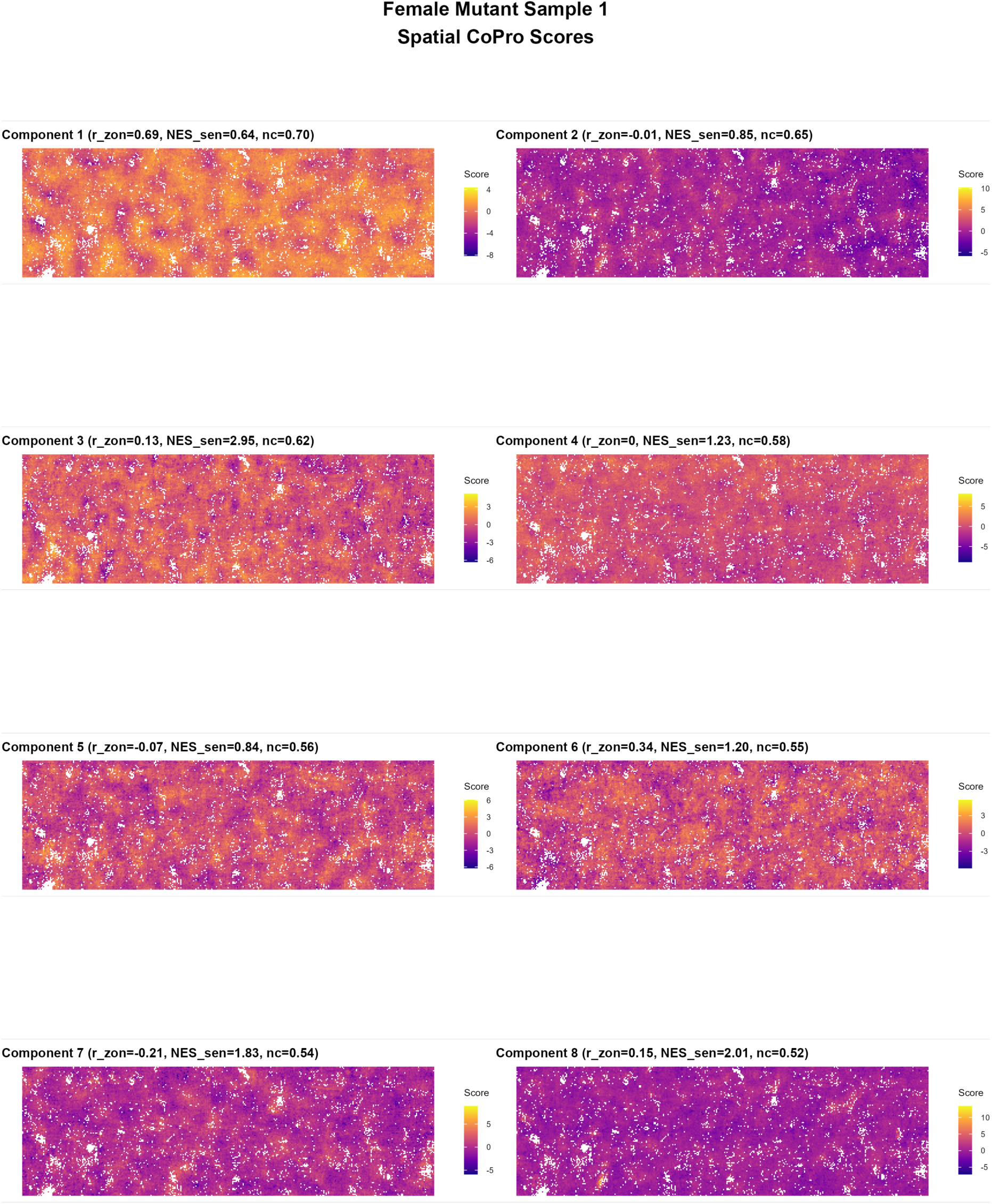
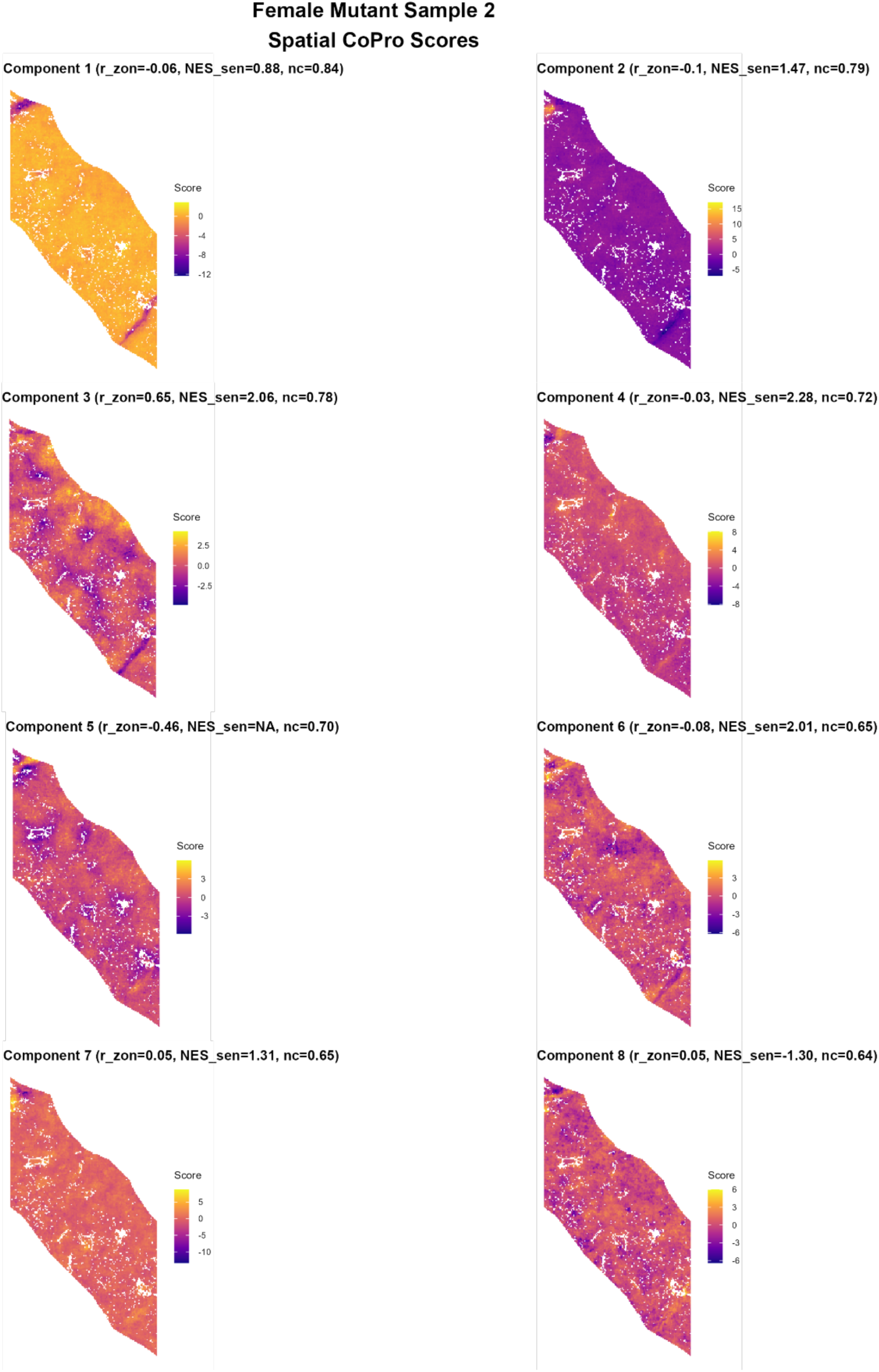
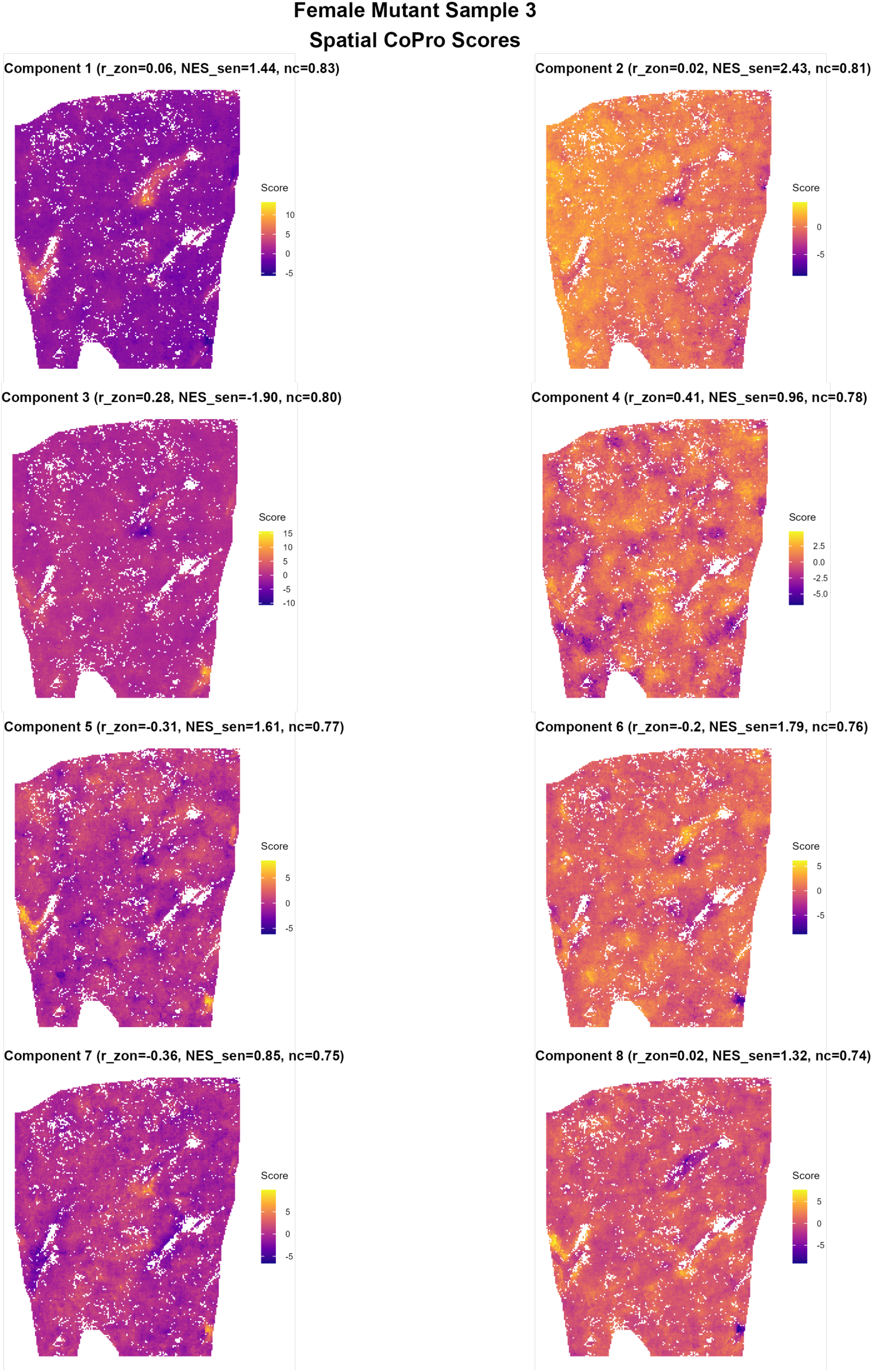
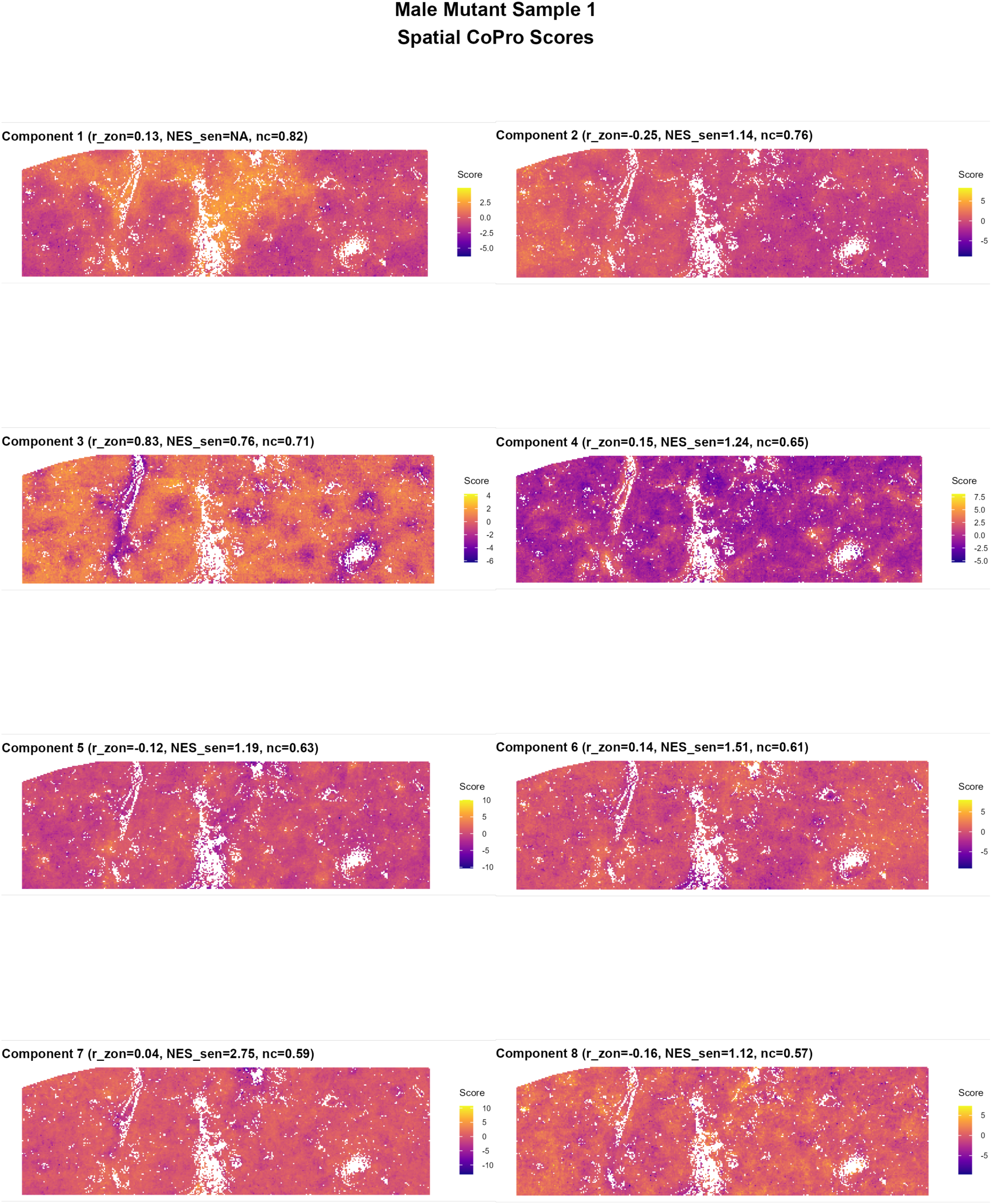
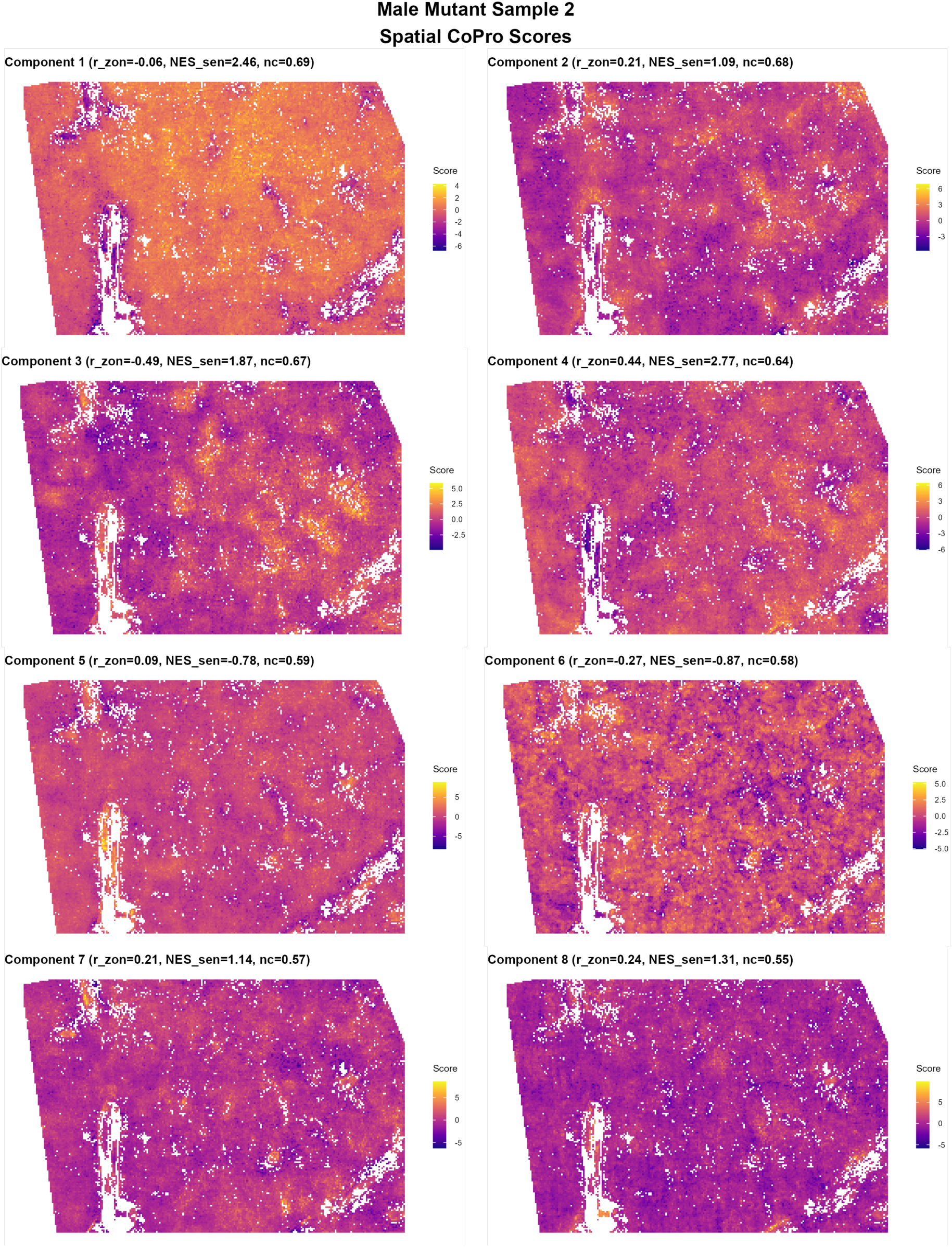
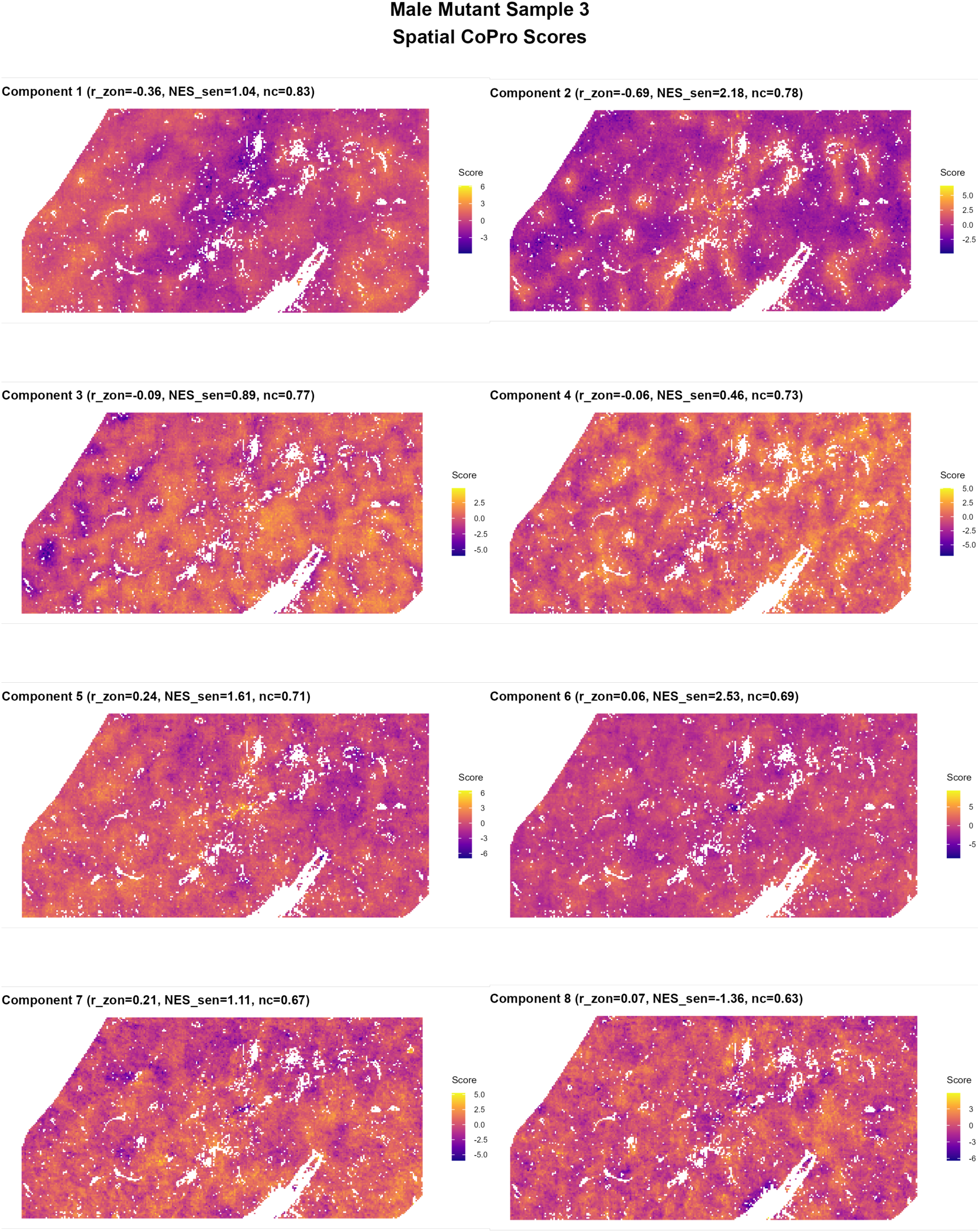
Spatial raster plots showing the cell score for each of the eight CoPro components identified in each of the mutant liver samples (female replicates 1-3, followed by male replicates 1-3) Each panel displays the spatial distribution of component scores across the tissue, with color indicating relative enrichment. Component titles indicate the correlation with zonation (r_zon), enrichment for the senescence gene set (NES_sen), and normalized component strength (nc). White regions correspond to vascular structures where no expression measurements were obtained.

**Supplementary Figure 13.**
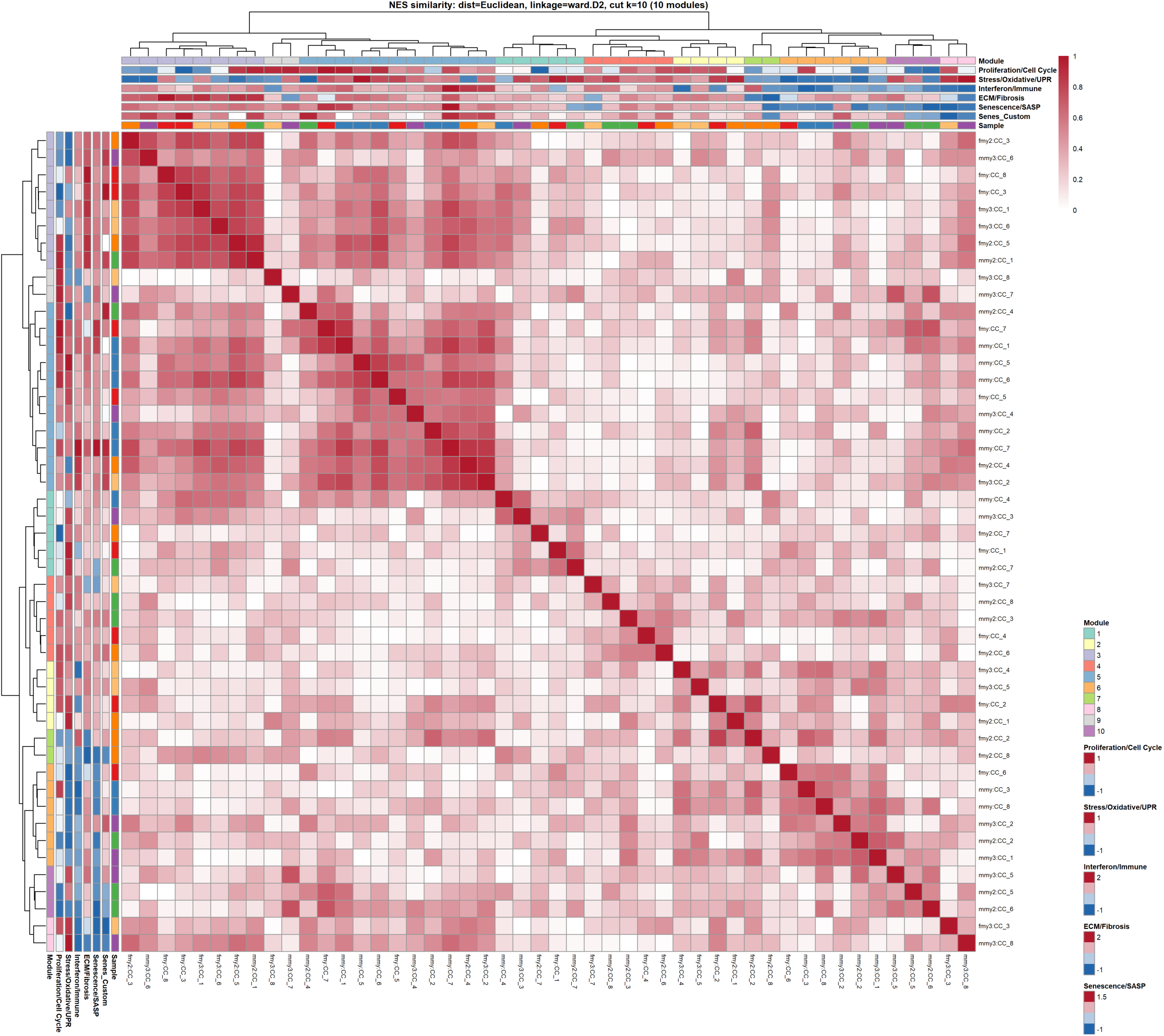
Heatmap showing pairwise similarity between all 48 CoPro components (six samples × eight components per sample), computed from Euclidean distances of normalized pathway enrichment scores (NES) and clustered using Ward.D2 linkage. Components with similar pathway signatures cluster into transcriptional modules (color annotation). Additional annotations summarize enrichment across pathway themes, including proliferation/cell cycle, stress/oxidative response, interferon/immune signaling, extracellular matrix (ECM)/fibrosis, and senescence/SASP programs. This analysis reveals reproducible aging modules shared across animals despite variation in their spatial localization.

**Supplementary Figure 14.**
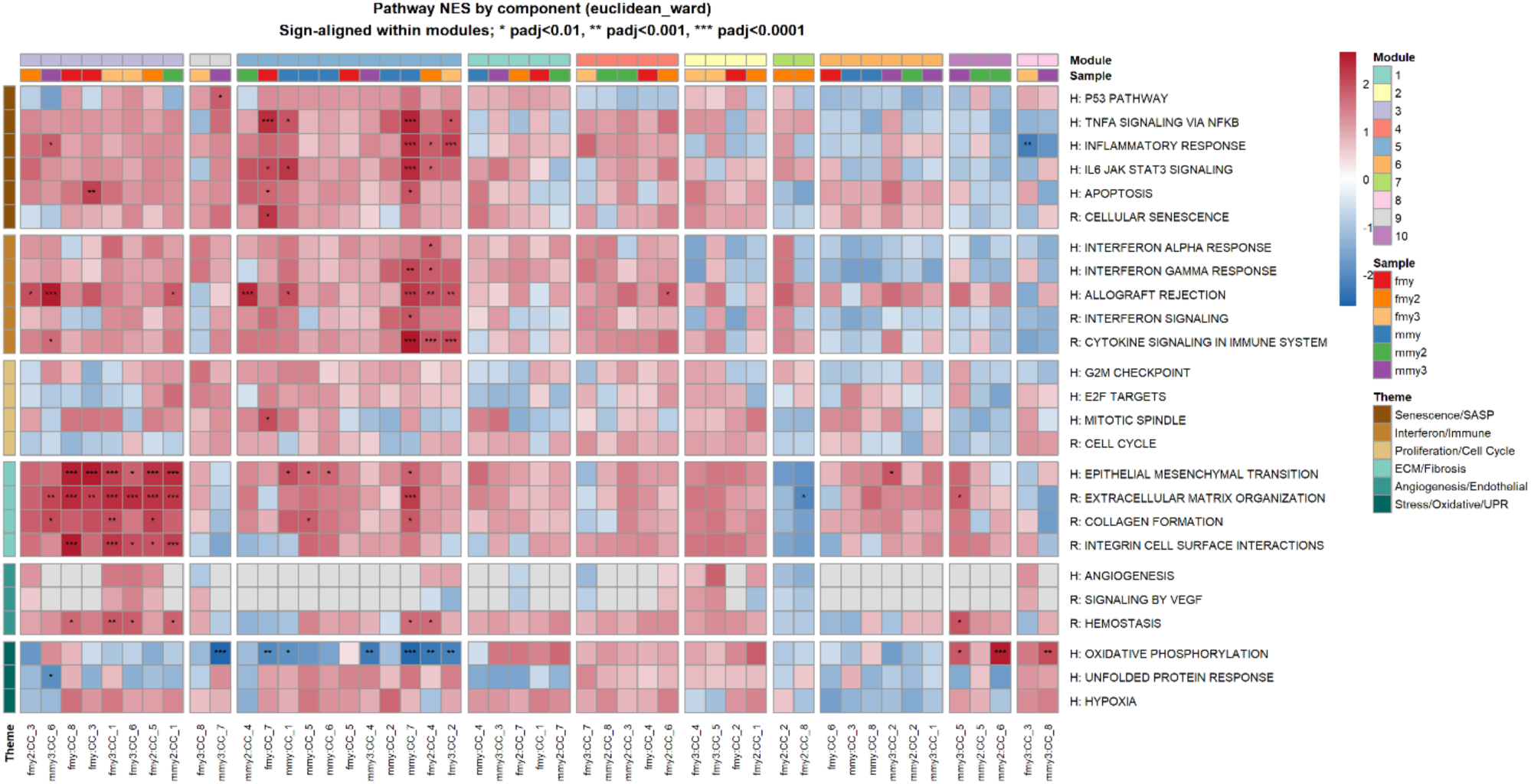
Heatmap showing normalized enrichment scores (NES) for pathway signatures associated with each CoPro component (six samples × eight components per sample). Columns correspond to components and rows to pathways grouped into biological themes including senescence/SASP, interferon/immune signaling, proliferation/cell cycle, ECM/fibrosis, angiogenesis/endothelial programs, and stress/oxidative responses. NES values were sign-aligned within modules to facilitate comparison across components. Components are grouped by hierarchical clustering based on Euclidean distance of pathway enrichment profiles, revealing reproducible transcriptional modules shared across animals. Asterisks indicate statistical significance of pathway enrichment.

**Supplementary Figure 15-17.**
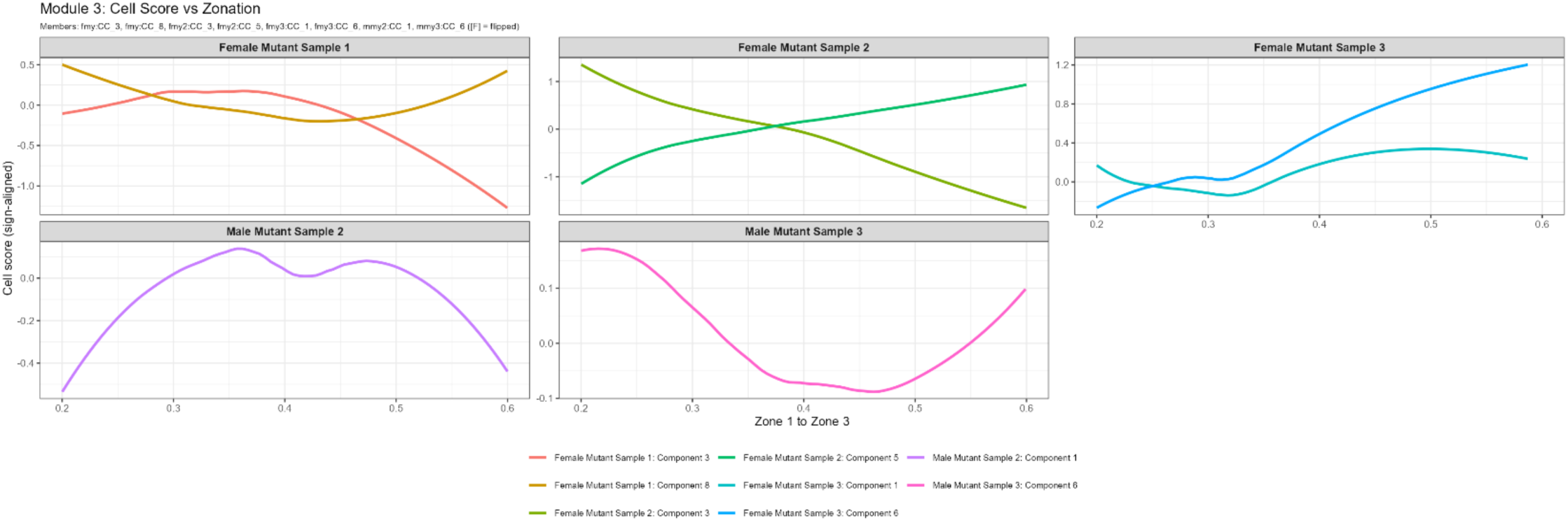
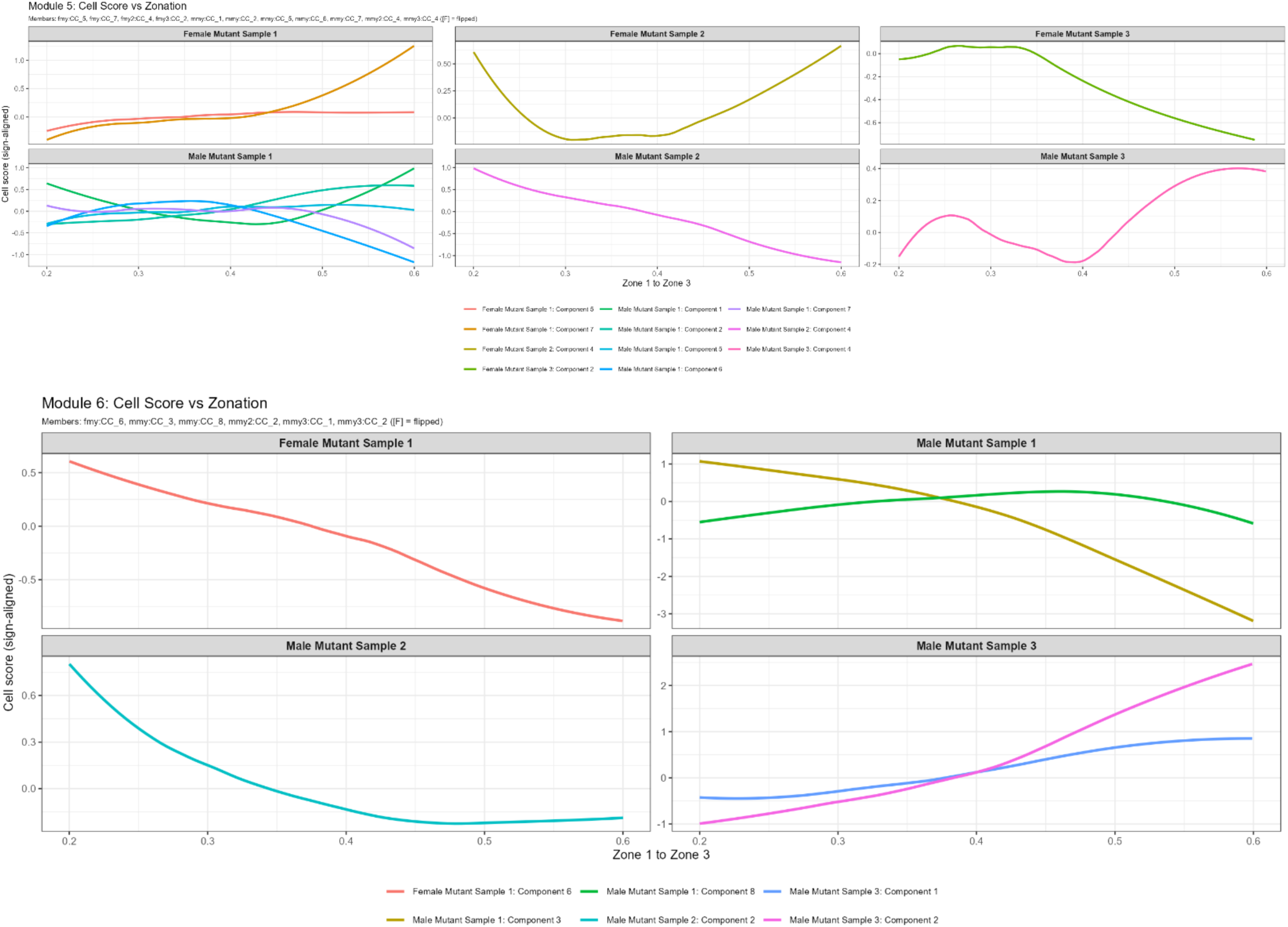
Relationship between Module component scores and lobular zonation for modules 3, 5, 6. Local spline-fits showing the relationship between CoPro component scores and the zonation coordinate for all components assigned to Module 3 across mutant liver samples. Each curve represents a component from an individual sample, with component scores sign-aligned to enable comparison.

**Supplementary Figure 18.**
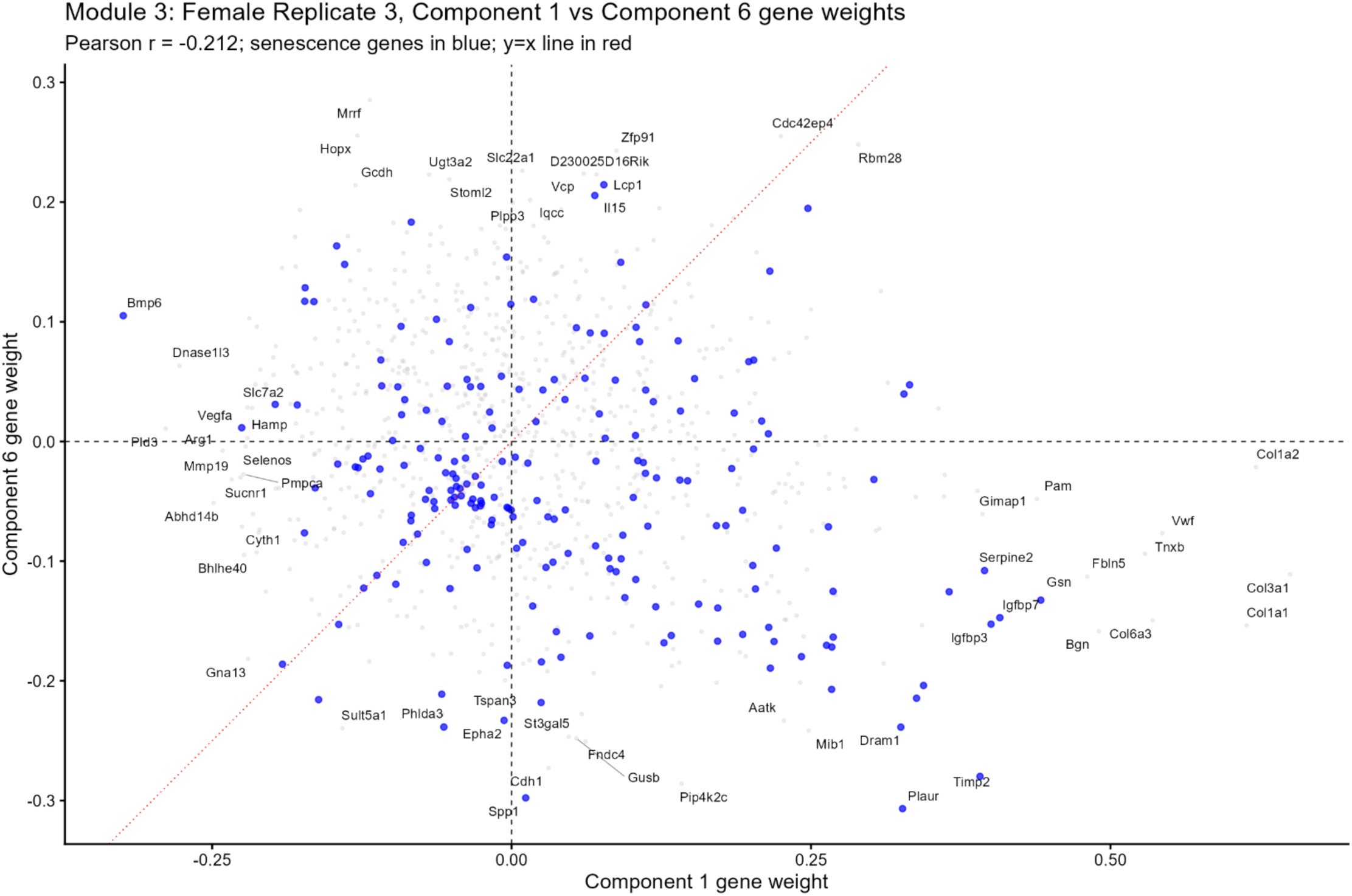
Gene weight comparison of two components within the same ECM-dominated module in female replicate 3. Scatter plot of gene weights for Component 1 (x-axis) versus Component 6 (y-axis), both members of Module 3 in the Euclidean-ward clustering of pathway enrichment profiles. Each point represents one gene; senescence-associated genes are highlighted in blue. The diagonal red line indicates y = x.

**Supplementary Figure 19.**
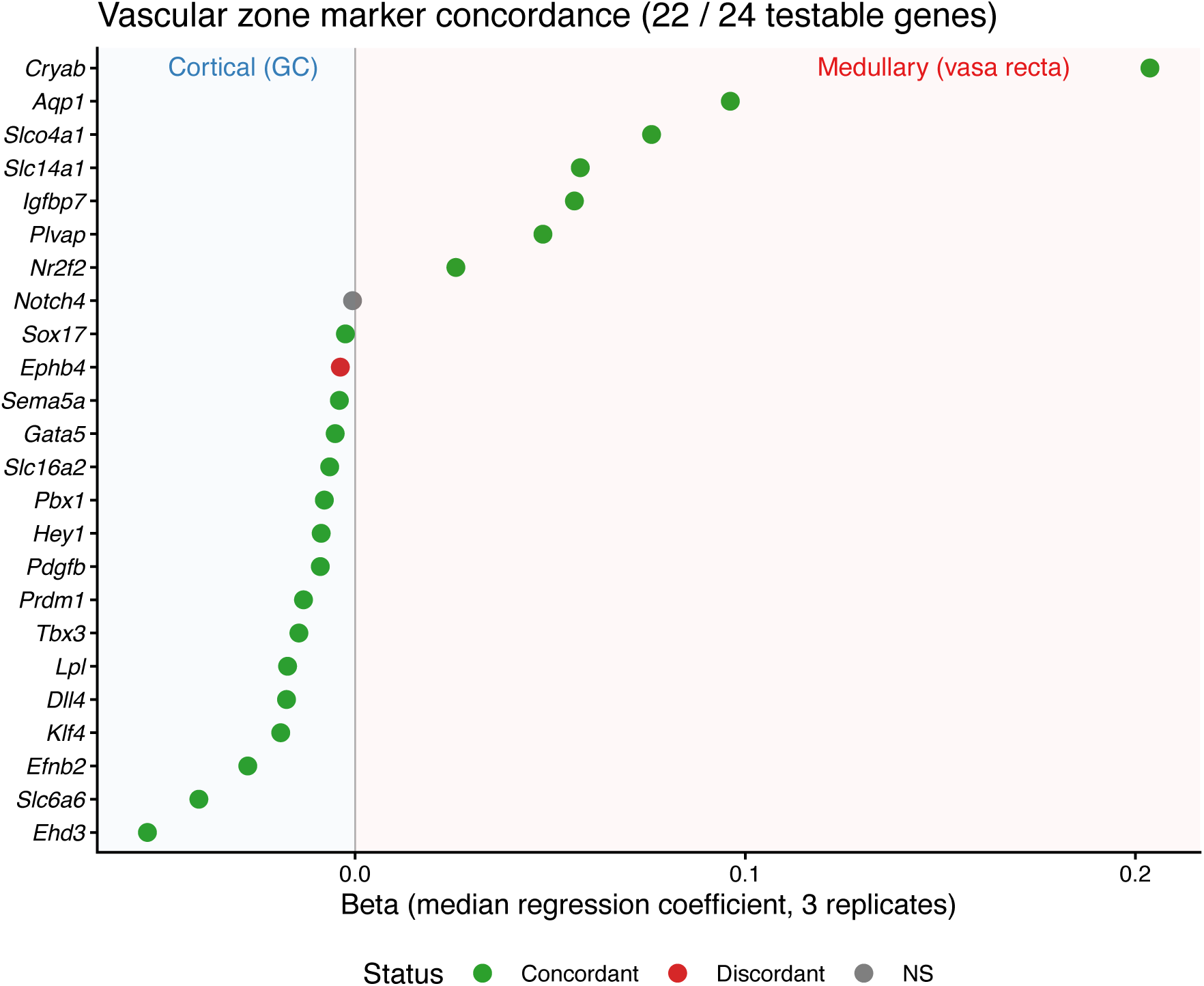
Thirty zone-specific vascular marker genes from Barry, et al. (2019) were evaluated against the CoPro-derived corticomedullary vascular axis. Pan-endothelial and non-zone-specific genes were excluded, leaving 24 testable genes (6 were filtered due to expression in <1% of cells). Each gene is plotted by its median regression coefficient (beta) across three replicates, ordered from cortical/glomerular capillary (negative) to medullary/vasa recta (positive). Points are colored by concordance status: concordant (green), discordant (red), or non-significant (grey).

